# Inhibition and excitation shape activity selection: effect of oscillations in a decision-making circuit

**DOI:** 10.1101/373282

**Authors:** Thomas Bose, Andreagiovanni Reina, James A.R. Marshall

## Abstract

Decision-making is a complex task and requires adaptive mechanisms that facilitate efficient behaviour. Here, we consider a neural circuit that guides the behaviour of an animal in ongoing binary choice tasks. We adopt an inhibition motif from neural network theory and propose a dynamical system characterized by nonlinear feedback, which links mechanism (the implementation of the neural circuit) and function (increasing reproductive value). A central inhibitory unit influences evidence-integrating excitatory units, which in our terms correspond to motivations competing for selection. We determine the parameter regime where the animal exhibits improved decision-making behaviour, and explain different behavioural outcomes by making the link between bifurcation analysis of the nonlinear neural circuit model and decision-making performance. We find that the animal performs best if it tunes internal parameters of the neural circuit in accordance with the underlying bifurcation structure. In particular, variation of inhibition strength and excitation-over-inhibition ratio have a crucial effect on the decision outcome, by allowing the animal to break decision deadlock and to enter an oscillatory phase that describes its internal motivational state. Our findings indicate that this oscillatory phase may improve the overall performance of the animal in an ongoing foraging task. Our results underpin the importance of an integrated functional and mechanistic study of animal activity selection.

**Author summary:** Organisms frequently select activities, which relate to economic, social and perceptual decision-making problems. The choices made may have substantial impact on their lives. In foraging decisions, for example, animals aim at reaching a target intake of nutrients; it is generally believed that a balanced diet improves reproductive success, yet little is known about the underlying mechanisms that integrate nutritional needs within the brain. In our study, we address this coupling between physiological states and a decision-making circuit in the context of foraging decisions. We consider a model animal that has the drive to eat or drink. The motivation to select and perform one of these activities (i.e. eating or drinking), is processed in artificial neuronal units that have access to information on how hungry and thirsty the animal is at the point it makes the decision. We show that inhibitory and excitatory mechanisms in the neural circuit shape ongoing binary decisions, and we reveal under which conditions oscillating motivations may improve the overall performance of the animal. Our results indicate that inefficient or pathological decision-making may originate from suboptimal modulation of excitation and inhibition in the neurobiological network.

## Introduction

Modelling complex decision-making problems requires the integration of mechanism and function in a combined modelling framework [1–3]. In binary decision-making, for example, mechanistic neuro-models have been proposed that are based on different inhibitory motifs, such as cross-inhibition [4–10], feed-forward inhibition [11, 12], and interneuronal inhibition [13]. Furthermore, linear diffusion-type models which describe the accumulation of noisy evidence are widely applied to study two-alternative choice tasks [14, 15]. A specific feature of diffusion models is their ability to resemble statistically optimal processes [15]. However, it has been found that decision-makers do not always behave optimally [2, 8, 16–18]. Suboptimality may arise from the evolution of decision rules in complex environments [2, 16], from limited precision in neural computations [18], or from nonlinearity in the neural network circuitry, as nonlinear models often have several stable stationary states and therefore may integrate evidence by reaching a decision state, which may not correspond to the optimal solution [8].

The aim of the present paper is to study the behaviour of a hypothetical animal performing ongoing activity selection through a nonlinear neural circuit model that implements the interneuronal inhibitory motif. Unlike the cross-inhibition motif, in our model (and in the interneuronal inhibition motif in general [13]) evidence-integrating units do not inhibit each other mutually but receive negative feedback from a separate neural population - the inhibitory unit. Previous studies indicate that the cross-inhibition motif is an approximation of the more realistic interneuronal inhibition motif [7, 15] and we emphasise in the present paper that reducing model complexity may lead to the loss of dynamical regimes which show interesting and unexpected phenomena. In particular, we highlight that, compared with the cross-inhibition motif, our implementation of the interneuronal inhibition motif yields an extended subset of dynamical states characterising the decision-maker, including the occurrence of oscillations. In addition to the inhibitory motif, recurrent excitation is taken into account in the same decision-making circuit. The excitation-over-inhibition ratio (E/I ratio) plays a pivotal role in our model analysis and it is considered to be tunable in the decision-making circuit. In general, balancing excitation and inhibition in neural networks is crucial for processing input and executing functions [19–24], and modulating excitation and inhibition can drive the decision-maker through different states characterising the decision-making performance [25, 26]. Specifically, cognitive disorders are related to unbalanced E/I-ratios [27], which may be reflected by impaired decision-making present, for instance, in schizophrenia where deficits in inhibition of behavioural responses can be observed [28].

To examine the utility of our proposed mechanism we embed it in a well-studied nutritional theory, the geometric framework [29]. In this framework animals perform actions (consume resources with different nutrient ratios) to reach a preferred nutrient target. They derive utility according to how close to the target they get, usually determined by Euclidean distance between momentary nutritional state and target state, as in the present study. This framework, while capturing real feeding behaviour of diverse species [29], seems sufficiently general to capture the general problem of ongoing action selection [30]. More specifically, in the scenario considered in the present paper, a model animal makes ongoing foraging decisions and may perform the action chosen until another food choice is made. Our modelling approach is based on the assumption that, influenced by the level of nutrients inside the body, animals have an inner drive (or motivation) to feed and drink [31–34]. This nutritional state changes over time and therefore animals have to select and perform activities that satisfy and balance their nutritional needs [29]. Furthermore, in real scenarios animals are embedded in an uncertain environment and may be subject to frequent changes [16] and predation risk [35]. To include uncertainties in our model, we assume that during the process of selecting the next activity and whilst performing the action chosen, the ongoing decision-making task may be interrupted. All information available to the decision-maker need to be integrated together with the momentary nutritional requirements in a multi-faceted physiologically- and neurobiologically-wired network [36–40]. Hence, the intake of required nutrients affects the decision-making process by reducing the motivation to feed or drink [33, 34, 41]. In this regard, the internal nutritional state acts as excitatory input for the underlying neural circuit involved in the decision-making process. In neurobiological networks, excitatory inputs are usually balanced by inhibitory mechanisms [19], and it has been shown in previous theoretical studies of foraging animals that inhibitory mechanisms facilitate improved feeding behaviour [33, 41].

Our model-based analysis can be considered as an approach to link nutritional state and behaviour coupled through a decision-making circuit. As a result, we find a mapping between the nutritional deficits of the animal combined with an adaptive tuning of excitation and inhibition and the animal’s foraging behaviour. As the ongoing decision-making of the animal resembles repeated two-alternative choice tasks, we conclude that low-performance decision-making of the animal emerges from unbalanced E/I-ratios, thus indicating potential parallels between neural disorders and ineffective intake of nutrients.

## Results

### Temporal evolution of inputs and internal representations

The model animal in our study is required to continuously make decisions between two activities. In what follows, we refer to activity 1 as ‘eating food’, and to activity 2 as ‘drinking water’. We take into account a cost for switching between both activities. This cost is given as time constant *τ* which represents the time necessary to overcome the physical distance between food source and water source. More details are given in the *Methods* section. Initially, we place the animal exactly midway between food source and water source, as illustrated in Fig 1. First, we assume that deficits in food (*d*_1_) and water (*d*_2_) are equal, i.e. ∆*d* = *d*_1_ – *d*_2_ = 0; below we also consider unequal deficits ∆*d* > 0 in Section *’Dependence of expected penalty on initial deficits’*.

**Fig 1.**
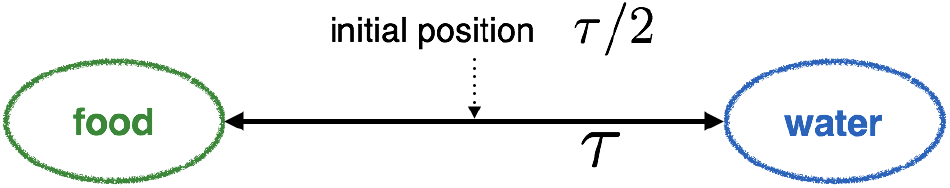
Depiction of the initial position of the animal. It is located at *τ*/2, i.e. exactly between food and water sources. The nutritional deficits in food and water are *d*_1_(*t* = 0) and *d*_2_(*t* = 0) when the ongoing decision-making process commences.

We start this analysis by presenting the results of a typical simulation. Fig 2 shows the evolution of food and water deficits (inputs) over time and the temporal evolution of the corresponding motivations (internal representations). We make the assumption that the animal decides to perform the activity with the greatest motivation. It feeds or moves towards the food source if ∆*x* > 0 and drinks or moves towards the water source if ∆*x* < 0. This assumption has been applied in previous studies of ongoing decision-making tasks, where internal representations are continuously updated over time (e.g. [41]). By inspecting Fig 2 we can see the interaction of physiological states (deficits) and neuronal variables (motivations). First, we allow the system to reach a stable stationary state (see Figs 2A and 2D) to ensure that we have well-defined initial conditions for the ongoing decision making task. Figs 2A and 2D show symmetric initial conditions, i.e. *x*_1_(*t* = 0) = *x*_2_(*t* = 0), with different absolute values that may be reached by oscillatory behaviour (Fig 2A). The presence of oscillations depends on the choice of the value of the E/I-ratio *r*. In addition, the E/I-ratio influences the motivations (Figs 2B and 2E) and the reduction of the deficits Figs 2C and 2F during the entire decision-making process. The *r* = 1 example shows that when the initial motivational state is close to dynamical regimes where oscillatory solutions are obtained (Fig 2A), then oscillations inherent to the nonlinear decision-making model also dominate during the ongoing decision-making task (Fig 2B). These oscillations around ∆*x* ≠ 0 are not present in Fig 2E, where the magnitudes |∆*x*| are generally smaller than in Fig 2B. Further to this, the *r* = 1 case is more effective than the *r* = 2 case, which we can see by comparing Figs 2C and 2F. For *r* = 1, the final state after the maximum bout time (see *Methods* section for definition of maximum bout time) is characterised by lower deficits compared with the *r* = 2 simulation. This finding indicates that oscillatory regimes which arise from nonlinearity of the underlying dynamical system may facilitate the continuous decision-making process.

**Fig 2.**
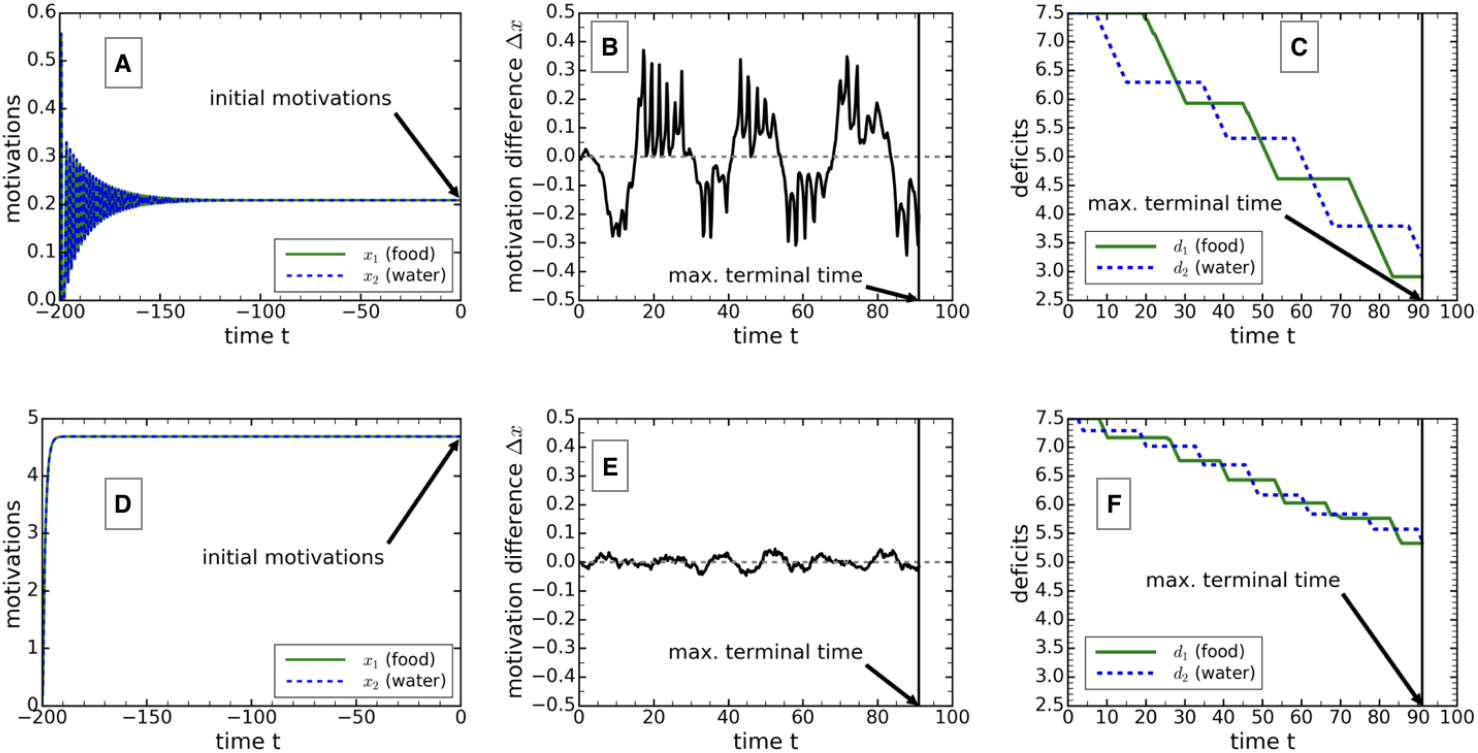
Simulation of ongoing decision-making process for different excitation/inhibition-ratios. A-C: *r* = 1 and D-F: *r* = 2. A and D show the evolution of the initial conditions for the motivations for *r* = 1 and *r* = 2, ending in well defined states. The animal is located at position *τ*/2 at time *t* = 0. The nutritional decision-making task starts at *t* = 0 and ends at *t* = 91, the maximum terminal time. The temporal changes of motivation differences are shown in B and E, with the corresponding reduction of deficits depicted in C and F, respectively. The values of the parameters are: *d_m_*(*t ≤* 0) = 7.5, ∆*d*(*t ≤* 0) = 0, *τ* = 4, *γ* = 0.15, *q* = 0.1, *β* = 3, *k* = 0.8, *k_inh_* = 0.8, *w* = 3, *g_e_* = 10 = *g_i_*, *b_e_* = 0.5 = *b_i_*, *σ* = 0 (A and D) and *σ* = 0.01 (B, C, E and F).

### Performance of the animal under the modulation of inhibition strength and excitation/inhibition-ratio

To further underline the results of the previous section, we present Fig 3, where we have simulated the system in Eq (4), introduced below in the *Methods* section, for inhibition strengths in the range 0 < *β* ≤ 5 and E/I-ratios varied between 0 < *r* ≤ 2.5. This graph depicts the performance of the hypothetical animal measured by the expected penalty (see Eq (5)), where lower values of the expected penalty correspond to a better performance. Additionally, we show the bifurcations that occur when varying the values of *β* and *r*. An area of improved performance is clearly recognisable (lowest values of expected penalty) in the bottom-left panel of Fig. 3. The shape of this area can be explained using the corresponding bifurcation diagrams. In Fig 3 we show the bifurcation diagram when keeping *β* = 3 constant and varying *r* (bottom-right panel), and the bifurcation diagram when keeping *r* = 1 constant and varying *β* (top-left panel). Both bifurcation diagrams relate to the initial deficit condition at *t* = 0. As time progresses, the bifurcation diagrams will be updated, so that at every instant in time the bifurcation diagrams change. However, we believe that the bifurcation diagrams at *t* = 0 are the most important ones and representative for the whole decision-making process, as the system has been prepared in one of the available stationary states. Furthermore, during the ongoing nutritional decision-making task, the animal goes through several internal changes, where deficits are equal (due to alternating motivations and intake of nutrients). In all these cases, the bifurcation diagrams show similarities. However, for *t* > 0 it is unlikely that the animal reaches one of the stable steady states available, as the decision-making process is subject to continuous updates of the deficits (whilst the animal feeds or drinks) and therefore frequent internal changes of nutritional deficit levels prevent the animal from reaching these states. This further indicates that the bifurcation diagrams at *t* = 0 are the most meaningful ones.

**Fig 3.**
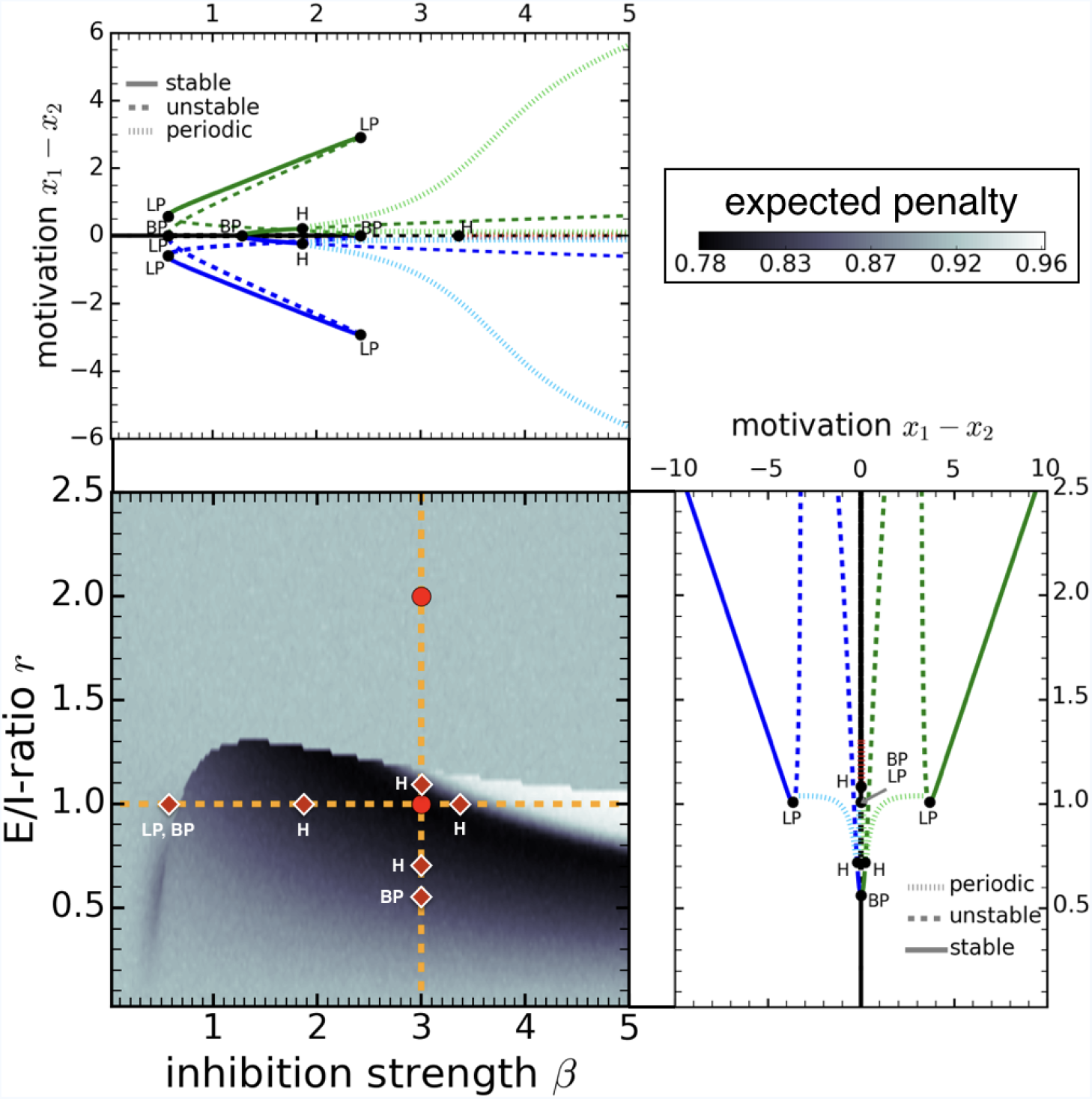
Depiction of the expected penalty with corresponding bifurcation diagrams. Bifurcation diagrams depending on inhibition strength *β* (top-left panel) and excitation/inhibition ratio *r* (bottom-right panel) correspond to orange dashed lines in bottom-left panel. Maximum and minimum amplitudes are plotted for the periodic solutions (top-left and bottom-right). Areas characterised by the lowest penalty values mirror the best performance of the model animal (bottom-left). The circular markers (bottom-left) correspond to the examples in Fig 2, i.e. the marker at (*β* = 3, *r* = 1) relates to Figs 2A-2C, and the one at (*β* = 3, *r* = 2) to Figs 2D-2F. The values of the parameters are: *d_m_*(*t* = 0) = 7.5, ∆*d*(*t* = 0) = 0, *τ* = 4, *γ* = 0.15, *q* = 0.1, *k* = 0.8, *k_inh_* = 0.8, *w* = 3, *g_e_* = 10 = *g_i_*, *b_e_* = 0.5 = *b_i_*, *σ* = 0.01 (bottom-left) and *σ* = 0 (top-left and bottom-right). Abbreviations: LP: limit point, BP: branch point, H: Hopf bifurcation point. Selected bifurcation points from the top-left and bottom-right panels are redrawn in the bottom-left panel as diamond markers along the orange dashed lines.

Inspecting the bifurcation diagram when *β* is the critical parameter (top-left panel), we can see that for low values of the inhibition strength (*β* < 0.57) the only stable fixed point is given by a decision deadlock state (∆*x* = 0). Increasing *β* to larger values, we observe possible decision deadlock-breaking indicated by the existence of stable equilibria with 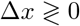. With the occurrence of decision deadlock-breaking the performance of the animal improves (compare bifurcation diagram in top-left panel and performance plot in bottom-left panel). The performance improves even more with the emergence of stable periodic orbits on additional branches 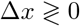 (see two Hopf bifurcations points for *β ≈* 1.9). However, at *β ≈* 3.4 we observe another Hopf bifurcation on the ∆*x* = 0 branch at which point the performance of the animal decreases (compare bifurcation diagram in top-left panel and performance plot in bottom-left panel).

Similar qualitative behaviour can be observed in the bifurcation digram with *r* as the critical parameter (see bottom-right panel). The performance improves as soon as the decision deadlock state is broken (see branch point at *r ≈* 0.56), and is even further enhanced with the emergence of stable periodic orbits on the 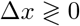 branches (see Hopf bifurcations at *r ≈* 0.71). However, we observe a clear drop in performance when the periodic orbits collide with saddle points and vanish (via homoclinic bifurcations at *r ≈* 1.0; homoclinic bifurcations are further discussed in Section 2 in S1 Text). In addition, at *r ≈* 1.1 we observe another Hopf bifurcation on the ∆*x* = 0 branch, which may contribute to the performance drop. Periodic limit cycles relating to these Hopf bifurcations, however, only exist until *r ≈* 1.3, where we observe a limit point of cycles. Here, stable and unstable periodic orbits meet. The unstable orbit vanishes at approximately the same *r*-value.

Our results indicate that the occurrence of periodic orbits on branches characterised by motivation differences 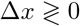 may enhance decision-making performance. The size of the area of improved performance is more extended along the *β*-axis and narrower along the *r*-axis, which seems to be strongly correlated with the range for which these periodic solutions exist. In contrast, periodic oscillations on the ∆*x* = 0 branch lead to a drop in performance. If the motivational state is moving along this orbit, then frequent changes in motivation difference may occur due to the presence of noise. The stable orbit, however, prevents the solution from gaining large motivation differences and drives it back to the symmetric state *x*_1_ = *x*_2_. In contrast, when the motivation move along the asymmetric orbits with 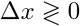 (e.g. see Fig 2B) the periodic orbit allows the motivations to achieve sufficiently large motivation differences, so that the animal can effectively perform one of the actions, feeding or drinking. However, within one cycle motivation differences always come close to the switching line ∆*x* = 0. Due to the reduction of deficits (whilst eating or drinking) and the presence of noise, this allows activity switching in an efficient way. As food and water sources are physically separated, oscillatory solutions seem to be a way to mediate between feeding and drinking behaviour of the animal determined by motivation differences computed in a neural circuit, the cost for switching (the animal has to travel without being able to reduce deficits) and the internal nutritional state.

#### Possible effects of noise on behaviour modulated by excitation and inhibition

Inside the brain, noise is present at all stages of the sensorimotor loop and has immediate behavioural consequences [42]. Varying noise strengths may induce transitions between different dynamical regimes [43–45]. For example, it has been shown that the presence of noise in nonlinear dynamical systems may shift Hopf-bifurcation points [43], and can lead to stochastic resonance-like behaviour even in the absence of external periodic signals, when the system is close to a Hopf-bifurcation point [44]. This seems to be particularly relevant for our study, as we have demonstrated that stable limit cycles born at Hopf-bifurcation points may improve decision-making and feeding behaviour. Considering the noise strength *σ* as the critical bifurcation parameter in our model (see Eq (4)), it will be interesting to see if the behavioural response of the model animal can be further improved by tuning *σ* within an appropriate parameter range. This, however, is a subtle issue and deserves to be investigated in a separate study, as noise-induced Hopf-bifurcation-type sequences may also arise in parameter regimes, where noise-free equations do not exhibit periodic solutions [45]. We consider this topic of behavioural resonance as a possible direction for future research.

### Dependence of expected penalty on initial deficits

To investigate the dependence of the expected penalty on the deficit difference at *t* = 0 we refer to Fig 4, where expected penalties are plotted for different E/I-ratios *r*, alongside examples of the temporal evolution of motivations for selected ∆*d*(*t* = 0). To investigate the effect of nonlinearity, we show the results for the nonlinear model in Eq (4) in Figs 4A-4C and for a linearised version of that model in Figs 4D-4F. The mathematical details of the linearisation are given in Section 5 in S1 Text. To simplify the comparison among different ∆*d*(0), we choose the initial deficits *d*_1_(0) and *d*_2_(0) such that the value of the initial penalty *p*_0_ remains constant for all ∆*d*(0). Hence, in all cases the animal’s deficit state is characterised by identical initial penalties but different initial deficits.

**Fig 4.**
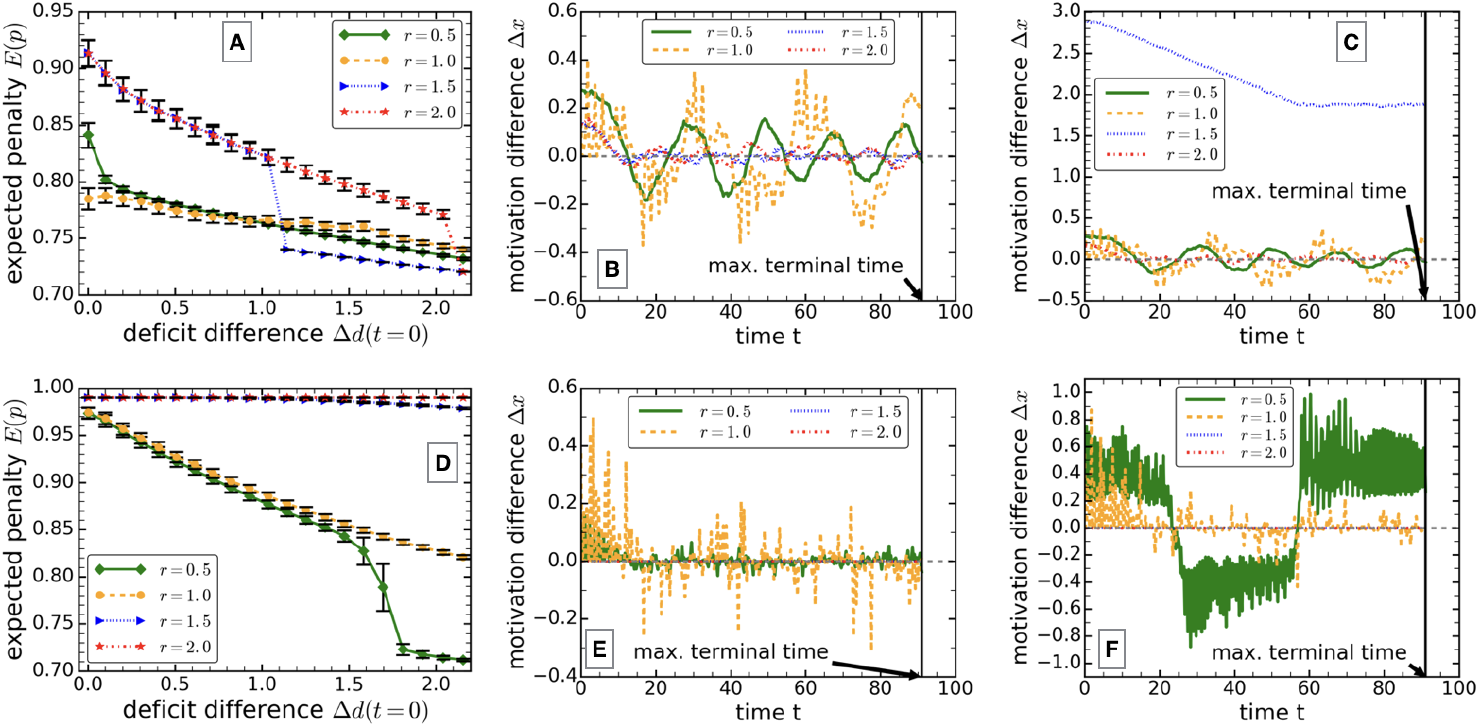
Plots of expected penalties and motivations depending on the initial deficit difference. Curves are shown for different E/I-ratios. In all curves the initial deficit difference was introduced such that the initial penalty was kept constant. This means that the mean value of the initial deficit was shifted to lower values with increasing ∆*d*, i.e. we have a range from *d_m_*(*t* = 0) = 7.50 for ∆*d*(*t* = 0) = 0 up to *d_m_*(*t* = 0) = 7.42 for ∆*d*(*t* = 0) = 2.16. A-C: nonlinear model in Eq (4); D-F: linearised model presented in Eq (S6) in S1 Text. Error bars in A and D represent standard deviations obtained from 1000 simulations. B: ∆*d*(0) = 1.04 (nonlin. model), C: ∆*d*(0) = 1.25 (nonlin. model), E: ∆*d*(0) = 1.25 (lin. model), F: ∆*d*(0) = 1.81 (lin. model). The values of the other parameters are: *τ* = 4, *β* = 3, *γ* = 0.15, *q* = 0.1, *k* = 0.8, *k_inh_* = 0.8, *w* = 3, *g_e_* = 10 = *g_i_*, *b_e_* = 0.5 = *b_i_*, and *σ* = 0.01.

Fig 4 shows the impact of varying the E/I-ratio *r* on the performance of the animal depending on the initial deficit difference. For instance, the *r* = 1 curve in Fig 4A shows a lower penalty value compared with both smaller (*r* = 0.5) and larger (*r* = 1.5 and *r* = 2) values of the E/I-ratio for sufficiently small differences in the initial deficits. In contrast, when increasing the initial deficit differences we can see that first the *r* = 0.5 and *r* = 1.5 curves (at ∆*d*(*t* = 0) ≈ 1) and later the *r* = 2 curve (at ∆*d*(*t* = 0) ≈ 2.1) drop below the *r* = 1 curve. However, the *r* = 0.5 and *r* = 1.0 show only small differences in performance in the whole ∆*d*-interval, except for very small ∆*d*(*t* = 0) (Fig 4A). Thus, we find that adjusting the E/I-ratio according to the initial deficit state may help the animal to improve its effectiveness. The drop of expected penalty we observe on the *r* = 1.5 and *r* = 2 curves in Fig 4A is a direct consequence of the interplay between switching cost *τ* and the coexistence of different stable stationary motivational states. The existence of these stationary states is characterised in detail in Section 2 in S1 Text and briefly described in the following. For ∆*d >* 0 there are two different stable fixed points available with ∆*x >* 0, one characterised by a large difference in motivations and another fixed point characterised by a small motivational difference. In what follows, the value of the initial deficit difference quantifying the switch between small-∆*x* and large-∆*x* stable fixed points is denoted ∆*d_switch_*. Consider, for example, the *r* = 1.5 curves in Fig 4A and 4B. If the initial deficit difference is small (0 ≤ ∆*d <* ∆*d_switch_ ≈* 1.0, Fig 4A), then motivational differences are small, too (see initial motivations for *r* = 1.5 at *t* = 0 in Fig 4B). However, if initial deficit differences are larger than ∆*d_switch_*, the initial motivational states make a transition from the small-∆*x* fixed point to the large-∆*x* equilibrium (cf. initial motivations for *r* = 1.5 at *t* = 0 in Figs 4B and 4C). If this occurs, then the motivational differences are so far away from the switching condition (∆*x* = 0) that the animal only consumes one item, either water or food, over the entire course of the ongoing decision-making task. Even when the animal has reduced all deficits of one type to zero, its motivations reach a new steady state which is still too far from the switching condition, as shown in Fig 4C (see curve labelled *r* = 1.5). The explanation for the drop of the *r* = 2 curve at ∆*d ≈* 2.1 in Fig 4A is equivalent to that for the behaviour of the *r* = 1.5 curve. Hence, for sufficiently large differences of the initial deficits the animal may consume only one nutritional item, and by doing so, may achieve the lowest penalty value. However, this is only beneficial if the corresponding time frame is sufficiently small. Otherwise, it would be detrimental for the animal to only focus on balancing one of its deficits, and neglecting the other one. We also note that for zero (or sufficiently small) switching costs, the penalty for consuming exclusively one item (here food or water) would be higher compared with switching between the two activities [41, 46]. We confirm this and present more details about the reduction of deficits for *τ* = 0.05 and *τ* = 4 in Section 7 in S1 Text, including the deficit plots corresponding to Fig 4C.

Results obtained based on the linearised interneuronal inhibition model are depicted in Figs 4D-4F. Here, we also observe improved performance with increasing initial deficit differences. In particular, we observe a performance boost on the *r* = 0.5 curve at ∆*d ≈* 1.7 (Fig 4D). The reason for the drop in penalty is that the animal enters oscillatory motivational states that are sufficiently far away from the switching line, but allow switching after appropriate deficit reduction and due to noise (compare *r* = 0.5 curves in Figs 4E and 4F). Hence, we observe qualitative similarities between the nonlinear model in Eq (4) and its linearised version (see Eq (S6) in S1 Text). Nevertheless, only in the nonlinear system do we observe the coexistence of several possible equilibria, whereas in the linear system there is only one such state available, which can however change with modulation of the E/I-ratio.

In Section 8 in S1 Text we also show that with increasing distance between food and water sources (i.e. switching cost *τ*), the expected penalty increases as well, making feeding and drinking less efficient for larger physical distances that have to be overcome to take in nutrients. Further to this, we also demonstrate that an equal increase of initial deficits accelerates the reduction of deficits in Section 9 in S1 Text. This is a clear indication that the interneuronal inhibition model employed in the presented paper is sensitive to input magnitudes, which has emerged to be a general feature of both individual [9, 47, 48] and collective decision-making [30, 49–51]. We present more details on magnitude-sensitivity of our interneuronal inhibition model in Section 1 in S1 Text. Finally, we point out that the expected penalty also depends on the deficit decay constant *γ*, which is discussed in Section 10 in S1 Text.

## Discussion

Using a realistic neural inhibition motif, we demonstrated that modulating inhibition and E/I-ratio may enhance decision-making performance in an ongoing binary decision-making task. The inhibitory motif was implemented in a simple coarse-grained circuit that focuses on mechanism without taking into account synaptic details. This approach has proven to be useful in previous studies, where models have been introduced as low-dimensional right from the start [4–6, 15] or have been derived via dimension reduction from more complex network models [7, 26, 52]. Critically, we find that the model can be driven through different states by the regulation of parameters of the underlying neural circuit, thereby controlling the efficiency of the decision-making task. This is in qualitative agreement with results obtained from neural network analysis [25], where it was found that independently modulating postsynaptic excitatory and inhibitory conductances may increase decision-making performance and robustness. However, a pathological relationship between excitation and inhibition may diminish neural network stability and hence decrease the signal-to-noise ratio, which, for example, could account for different symptoms in schizophrenia [53], a neural disorder that causes failures in cognitive control [28]. Hence, our model analysis further underlines that modulating excitation and inhibition is of paramount importance in the ability of neural circuits to adapt to change in a controlled way, in accordance with previous studies on the interplay of excitation and inhibition in neuronal activity [19–23].

We find that stable limit cycles, which emerge at Hopf bifurcation points, are accessible states of the decision-making circuit, which have not been reported previously in other low-dimensional decision-making models (cf. [4–7, 26], for example). The existence of stable limit cycles is influenced by the magnitude of excitatory inputs (Fig 4), and the adjustment of inhibition strength *β* and E/I-ratio *r* in response to the stimulus (Figs 2 and 3). Strikingly, our results show that oscillating internal representations of the decision variable may have opposite effects — improvement or decline of decision-making performance — which can be explained by the underlying bifurcation structure of the model (Fig 3). Adaptation of internal neural parameters to varying stimuli in our model therefore points to a dual role of oscillatory activity in decision-making circuits. More generally, in the cortex, healthy oscillatory behaviour may be led back to the synchronisation of populations of cortical neurons, where interneurons (as in our study) are thought to play a pivotal role [19, 20]. Particularly in a decision-making context, electrophysiological recordings showed frequency-specific gamma-oscillations in parietal and fronto-polar cortex [54]. However, oscillations in corticobasal ganglia circuits are also related to neuronal disorders like Parkinson’s disease [55]. Moreover, recent findings suggest that intrinsic cortical oscillators are impaired in schizophrenia, which is supposed to originate from a lack of phase locking in brain oscillations; see [19] and references therein. These examples indicate that there are both positive and negative effects of oscillatory activity on cognitive abilities, in line with the findings reported in the present paper (e.g. see Fig 3).

Without loss of generality, we applied the nonlinear interneuronal inhibitory circuit to guide a model animal making foraging decisions. Doing so allowed us to link neuronal mechanisms, physiological states and functional foraging behaviour in the context of an ongoing decision-making task. In particular, we have shown that allowing both the inhibition strength *β* and the E/I-ratio *r* to vary provides the hypothetical animal with the ability to shape its response according to nutritional deficit levels, which is reflected in the animal’s performance (Fig 4). Interestingly, a study where monkeys performed motion-discrimination tasks, knowing about possible rewards associated with each option [56], indicates that animals can approach optimality in foraging decisions. In the present paper, the behaving model animal was embedded in an uncertain environment (interruption probability, e.g. predators might be present), and motivations were driven by nutritional imbalance (coupling of physiological and cognitive states). In our opinion, these assumptions together with calculating the animal’s performance using the distance from a target intake (geometrical framework [29]), enhance the comparability with foraging behaviour of animals.

Experimentally, it has been shown that predatory ground beetles (*A. dorsalis*), for instance, can regulate their intake of proteins and lipids [57], and that total egg production of female beetles of the same species peaked at the target intake of these two required nutrients [58]. This directly confirms the relationship between an optimal diet and improved reproductive success [58]. Similar results have recently been confirmed in mealworm beetles (*Tenebrio molitor L.*) [59], which further underline the immediate relationship between lifespan, reproductive value and a balanced diet [29, 46]. We note, that the ability to optimally adjust diets to nutritional needs has also been observed in decentralised systems, such as the monomorphic green-headed ants (*Rhytidoponera sp.*) [60], where social interactions may influence decision-making [61–63]. We believe that the implementation of the neural circuit linked with nutritional needs as investigated in the present paper is based on realistic assumptions, and therefore the model animal’s improved behavioural performance could be understood by balancing excitation and inhibition to enable an optimal diet, which directly relates mechanism with function. However, experimental studies have also revealed that decision-makers do not always act optimally [2, 15–18]. As noted before in [8], nonlinear dynamical systems may explain deviations from optimal behaviour as a result of different accessible states, that capture the decision-maker in suboptimal stable equilibria. Regarding our results, Fig 3 shows a separation between areas of good and bad performances, depending on the choice of inhibition strength *β* and E/I-ratio *r*. As underpinned by the corresponding bifurcation analysis, these distinct areas represent parameter regimes which offer different stable attracting motivational states. Therefore, improved (or optimal) foraging behaviour clearly depends on the optimal modulation of excitation and inhibition in the neural circuit, or in other words: a decline in performance is caused by deficits in the adjustment of excitation and inhibition.

In conclusion, our results may have important implications in understanding neuronal regulation in decision-making in general, and in a foraging context in particular. On the one hand, our findings may help elucidate the relationship between decision-making performance in ongoing decision-making tasks and impaired decision-making, and, on the other hand, may improve our understanding of the required adaptation of excitation and inhibition to appropriately modulate behavioural responses according to internal physiological needs and external factors.

## Methods

### The decision-making task

We consider a hypothetical animal which performs an activity-selection task whilst interacting with its environment. Driven by nutrional deficits, the task consists of deciding between two options – feeding or drinking. We take into account that the animal may be interrupted whilst executing the sequence of feeding and drinking bouts. This interruption might be due to the presence of a predator or fast changing weather conditions, for instance. To include the possibility of interruption, here we follow the modelling approach presented in [41] by assuming that feeding and drinking activities are geometrically distributed. The geometric distribution is given as
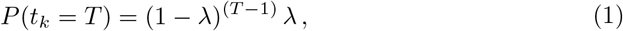

where *t_k_* takes integer values, i.e. *t_k_* = 1, 2, 3 With interruption probability λ the distribution *P* (*t_k_* = *T*) gives the probability that the ongoing decision-making task comes to an end at the integer time *t_k_* = *T*. In Fig 5 the geometric distribution and its cumulative distribution function (inset) are displayed. The maximum bout time *T_max_* is chosen such that at least 99% of the distribution is included.

**Fig 5.**
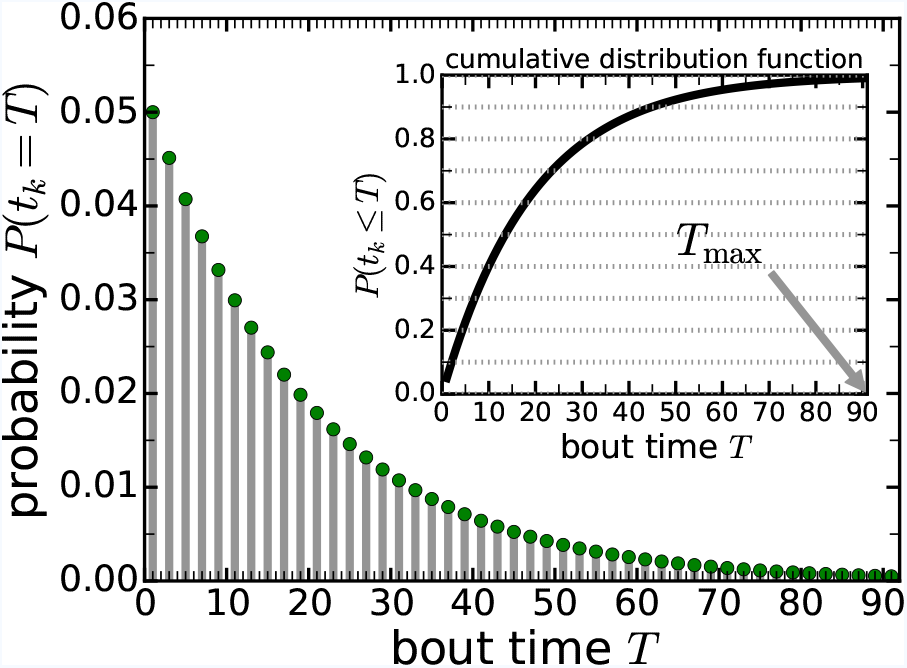
Geometric distribution. Plot of Eq (1) with interruption probability λ = 0.05. Interruption may happen at each integer time step. The inset depicts the corresponding cumulative distribution function.

In addition, we assume that food and water sources are physically separated. Thus, there is a cost for the animal to switch between feeding and drinking, as the animal cannot consume nutrients whilst it is moving to the place where it can perform the alternative activity. In our model (and as in the study in [41]) this cost is represented by a time constant *τ* quantifying the physical distance between food and water sources. Hereby we implicitly presume that the animal moves with an average speed, such that the linear relationship between position and time holds.

### Mechanism

To guide the behaviour of the animal making foraging decisions we adopt a model architecture that is based on a cortical network [13]. In particular, we use a nonlinear version of that model architecture, which has previously been studied as a simplified linear version [15]. A schematic of the model architecture is shown in Fig 6. The model relates to a decision between two food options. Evidence in favour of option 1 (option 2) is integrated by neuronal units *x*_1_ (*x*_2_). In nutritional decision-making option 1 may be ‘eat food’ and option 2 may be ‘drink water’. Choosing feeding and drinking is not a necessary assumption, as both options could represent any two food items that differ in their nutritional content, e.g. one food item rich in carbohydrates and the other rich in proteins, or, more generally, any two alternatives the magnitudes of which may vary over time. The momentary nutritional state of the decision-maker generates a representation as neural activation. This is the function of the pre-processing units in Fig 6, which transform the physiological state into inputs *I*_1_ and *I*_2_ that feed their respective integrators *x*_1_ and *x*_2_. Here, we assume a linear relationship between physiological levels characterising the nutritional state (deficits) and their representations in the neural circuit, i.e.

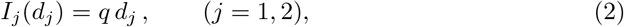

where *q* denotes the sensitivity of the animal to deficits *d_j_* (*j* = 1, 2) in nutritional items.

**Fig 6.**
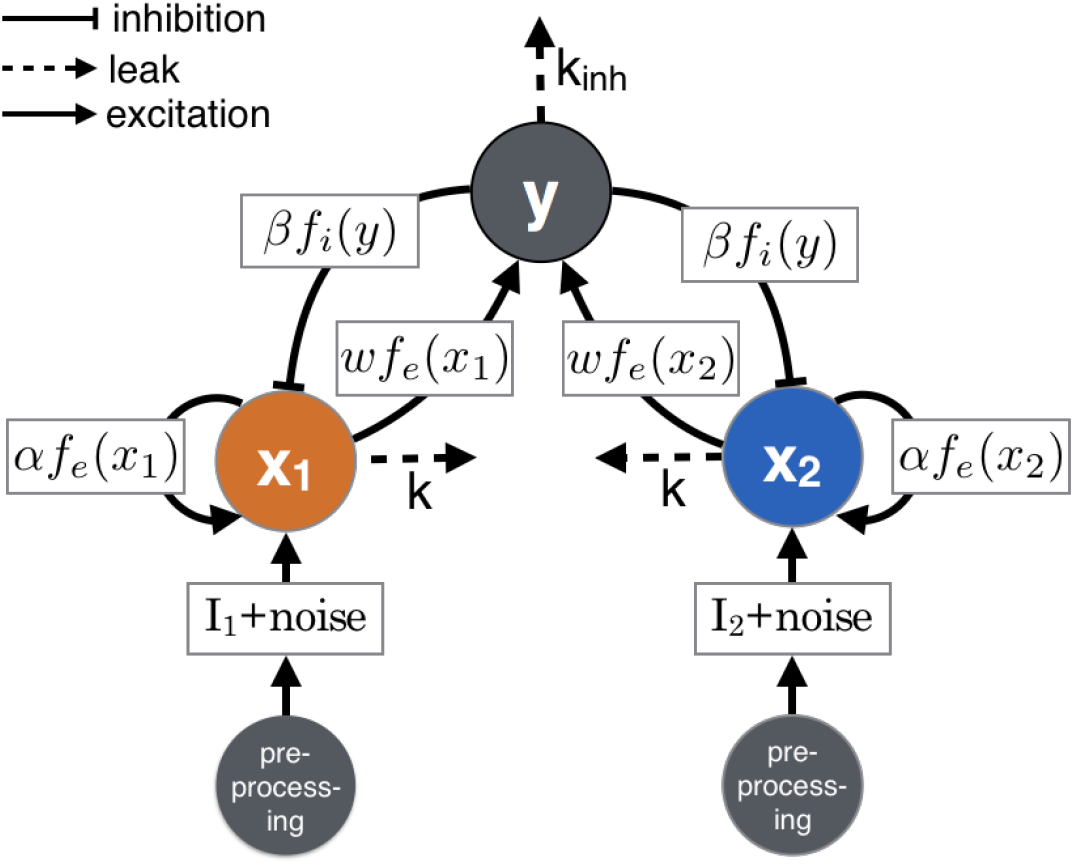
Schematic of the interneuronal inhibition motif. This is a graphical representation of the model in Eq (4).

Equation (2) corresponds to the curve labelled *g* = 2*/b* in Fig 7. As can be seen in this graphic, this curve, although based on a nonlinear functional dependency (of the same form as Eq (3)), approximately shows a linear relationship over a wide range of input values. We show in more detail in Section 4 in S1 Text under which conditions this assumption holds by approximating the sigmoidal relationship with a linear function, yielding a direct proportionality between *q* in Eq (2) and the gain characterising the sigmoidal function.

**Fig 7.**
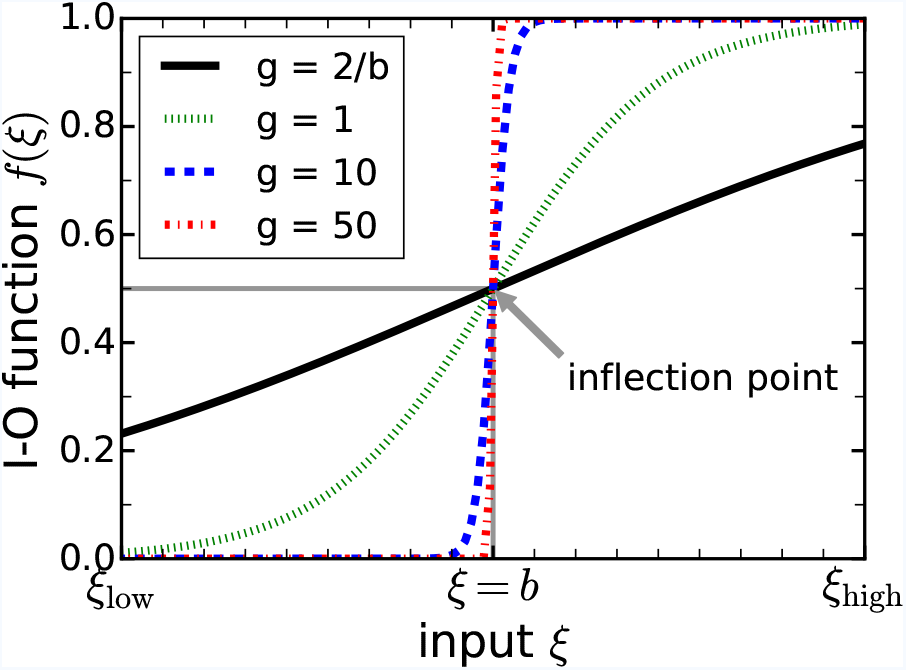
Plot of nonlinear input-output function. Typical input-output function that illustrates how neuronal units transform inputs via a nonlinear, sigmoidal function.

Further to this, we assume that inputs *I_j_* (*j* = 1, 2) are polluted by processing noise with standard deviation *σ*, which may arise from currents originating from other circuits in the brain. Noise is included via Wiener processes *W*_1_ and *W*_2_. Recurrent excitation is taken into account in the self-excitatory terms with strength *α*. If activity levels of *x*_1_ and *x*_2_ are sufficiently large then the interneuronal inhibitory unit *y* becomes activated with strength *w* and in turn inhibits the evidence-integrating units with strength *β*. The functions *f_e,i_*(⋅) appearing in different places in Fig 6 are nonlinear input-output (I-O) functions with a typical sigmoidal shape, as shown Fig 7. Here, *f_e_* denotes the nonlinear I-O function involved in excitatory processes, whereas *f_i_* represents inhibition. The form of the sigmoidal functions is given as

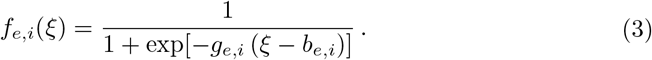

Here, *g_e_* and *g_i_* are the gains, and *b_e_* and *b_i_* are the positions where *f_e_* and *f_i_* have inflection points and reach half-level, respectively. In addition, we assume that information may be lost by including leak-terms in the evidence-integrating units *x*_1_ and *x*_2_ (rate *k*), and in the interneuronal inhibitory unit *y* (rate *k_inh_*). This model is described by the following system of nonlinear stochastic differential equations

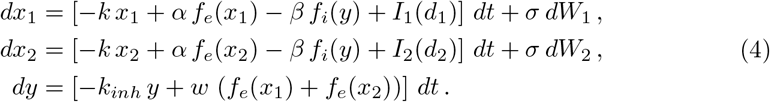

In addition to the nonlinear functions in Eq (4) we introduce an artificial nonlinearity to prevent *x*_1_, *x*_2_ and *y* from taking negative values, i.e. when numerically integrating the stochastic system from time *t_n_* to obtain the state variables at the next time step *t_n_*_+1_, we reset *X*_*t*_*n*+1__ = *max*(0, *X*_*t*_*n*+1__), *X* = *x*_1_*, x*_2_*, y*. Otherwise the leak terms (rates *k* and *k_inh_*) would become positive when the state variables change sign.

A detailed characterisation of this dynamical system is given in Section 2 in S1 Text. In the following, we identify *x*_1_ as the motivation to feed and *x*_2_ as the motivation to drink, and hence *d*_1_ and *d*_2_ represent food deficit and water deficit, respectively. During feeding and drinking, the time-dependent deficits are reduced according to *d_j_*(*t*) = *d_j_*(0) – *γ t*, where *d_j_*(0) are the initial deficits at *t* = 0 and *γ* is the deficit decay parameter [34, 41]. This is a valid assumption if feeding and drinking takes place within sufficiently short periods of time [34, 41].

### Relation with other models of decision-making

The model presented in Eq (4) is closely related to previous models of both perceptual and nutritional (value-based) decision-making. Therefore, it may be applied to a variety of decision-making problems, and may be incorporated into a hierarchy of models as follows. Consider the interneuronal inhibition model in Eq (4) detached from the foraging context. Although implemented as a nonlinear model in the present study, our model is a simpler, more tractable, version of the original network model proposed in [13], which included synaptic details and comprised 8000 equations. However, the pooled inhibition model investigated in [13] could be reduced in dimensionality to a model consisting of only two equations [7]. In terms of model complexity, our model can be considered as being in between the original model by [13] and the one which is reduced in dimension [7]. In Section 3 in S1 Text we explain that the model used in our paper offers additional dynamics (e.g. stable limit cycles) not present in the type of model studied by [7], which is based on cross-inhibition as inhibitory motif. The cross-inhibition motif has been applied in a variety of studies of decision-making [4–9] and may generally be considered as an approximation of the more realistic interneuronal inhibition motif [7].

To relate the model of the present paper to previous models of animal foraging behaviour [34, 41], we make use of a linearised version of the interneuronal inhibition model in Eq (4), which is derived in Eq (S6) in Section 5 in S1 Text, and derive from this linearised interneuronal inhibition model a model, which is based on the cross-inhibition motif in a linear two-dimensional dynamical system, see Eq (S8) in S1 Text. This approximated cross-inhibition model can be cast in equations, which closely resemble the model studied in [41], and under specific assumptions can be made equivalent to it. The mathematical details of this derivation are presented in Section 6 in S1 Text. As discussed in [41], the linearised cross-inhibition model can, in turn, be related to other models which describe animals making foraging decisions [34, 46] and models of perceptual decision-making [4, 15]. We note, however, that in contrast to previous models of activity selection (e.g. [34, 41]) the inputs in our model in Eq (4) do not contain derivatives *d*̇*_j_* (*t*), i.e. the rates of change by which deficits are altered over time. Given our model assumption that *d*̇_*j*_ = *−γ*, where *γ* is a constant, including them could be achieved by substituting *d_j_ → d_j_* + *c d*̇_*j*_ = *d_j_ – c γ*, where *c* is another constant (see also Section 6 in S1 Text). However, this might not hold for more complex physiological models, where the equation of motion describing the time evolution of the deficits *d_j_*(*t*) cannot be expressed in simple terms.

### Function

We adopt the assumption that in order to maximise its reproductive value a foraging animal needs to reach its target intake of nutrients [46]. Being empirically well motivated [29], we can further assume that reproductive value decreases with decline of the foraging performance, that is the distance between actual intake of nutrients and the target consumption increases [46]. A simple measure for performance in a nutritional decision-making task is the square of the Euclidian distance between current state and target state in nutrient space. Considering (super)organisms in nutrient space is known as the geometric framework and was pioneered by Simpson and Raubenheimer [29]. Several experimental studies have successfully confirmed the assumptions of this approach, e.g. see [58–60, 64–68].

In Fig 8 we show an illustration of the geometric framework in deficit space, where we assume that an animal has to decide about the sequence in which to feed or drink. Starting from an initial nutritional state characterised by deficits in water and food (state A), the animal selects the order of possible activities and how long it performs the activity chosen. In this example the animal switches several times between eating and drinking to reach the nutritional state B. The trajectory of the animal in Fig 8 is given as a combination of lines each of which is parallel to either the food or water deficit axis. This means that in our model the animal exclusively drinks or eats but does not perform both actions at the same time. This is further discussed below.

**Fig 8.**
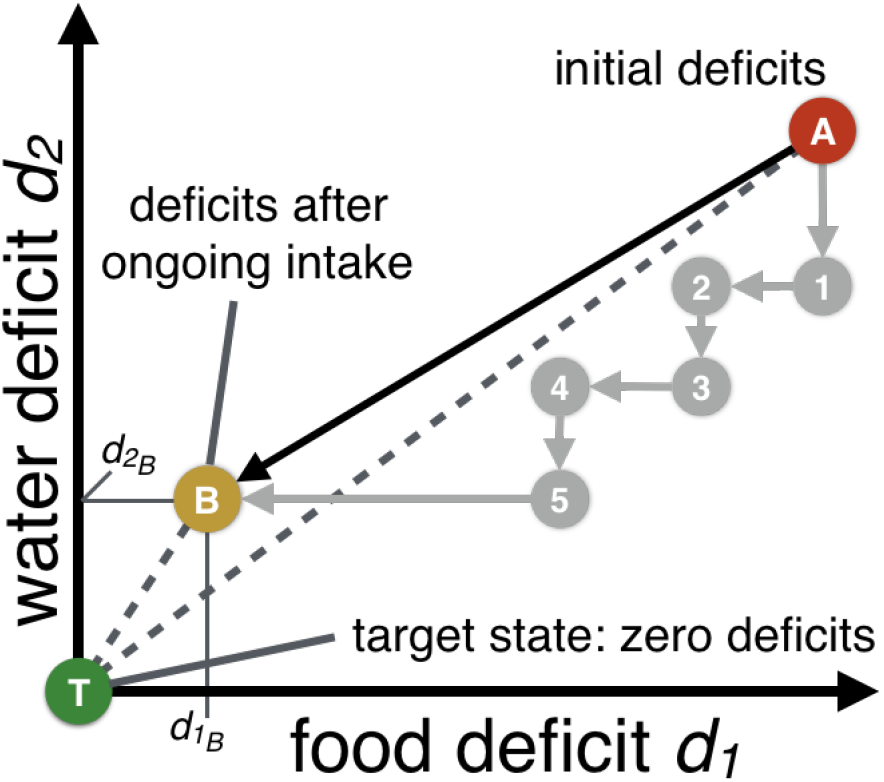
Illustration of the geometric framework. Application of the geometric framework of nutrition to deficit space; horizontal and vertical axes show the deficits of the animal. The target state is the origin of the diagram (*d*_1_ = 0, *d*_2_ = 0). Via a sequence of feeding and drinking bouts, the animal approaches its target state. Here, deficits can only decrease or remain constant. The performance is proportional to the Euclidean distance between states in the diagram and the origin, i.e. the target state T.

To characterise the momentary nutritional state and to evaluate behavioural performance, we make use of the penalty function 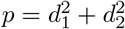, i.e. the square of the distance between points in the deficit space and its origin. For example, the penalty characterising state B is 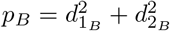. This penalty function has been applied in previous studies (e.g. [41]) and is equivalent to the reward function presented in [46]. Denoting the reward function *Rew*, then the relationship between reward *Rew* and penalty *p* is given as *p* = *K*_max_ – *Rew*, where *K*_max_ is a constant representing the maximum reward possible. Hence, a penalty of zero corresponds to yielding the maximal reward that relates to maximising reproductive value.

### Model implementation

The implementation of the neural circuit provides the link between the nutritional state of the animal and its foraging objective to come as close as possible to the target intake of nutrients. In every simulation, we place the animal initially exactly midway between food and water source. Initial deficits of food and water are set to either equal or unequal values. To determine the initial motivational state of the animal we use the dynamical system we employ in the decision-making process without noise (i.e. we set *σ* = 0 in Eq (4)) to obtain well defined initial conditions. This means that based on the interneuronal inhibition motif implemented in the neural circuit in Fig 6 the animal’s initial neural representation of the decision problem is also based on mechanism. This is different from previous theoretical studies (for example, cf. [41]), where an arbitrary initial motivational state has been assumed. During the execution of the ongoing decision-making task, processing noise is present in the neural circuit, and hence equal deficits will not lead to a decision-deadlock, which is in contrast to [41].

When numerically integrating the deterministic equations (*σ* = 0) we make use of a fourth-order Runge-Kutta method and when simulating the stochastic differential equations (*σ* = 0.01) we apply a predictor-corrector method, where the deterministic part is calculated with second order of accuracy in timestep *dt* [69]. For both methods we used a timestep of *dt* = 0.005 in the numerical integration. We found that this choice of *dt* gives a good compromise between computation time and accuracy when integrating the system, particularly with regard to the stochastic equations.

We interpret the activities *x*_1_ and *x*_2_ as motivations. The animal performs the activity with the greatest motivation, i.e. if *x*_1_ *> x*_2_ the animal chooses feeding and if *x*_2_ *> x*_1_ the animals selects drinking. However, the animal has to move to reach food source or water source and whilst the animal is moving we allow the motivations to change but the nutritional state is assumed to remain constant. Given the nonlinearity of the model, the inclusion of fluctuations and the continuous update of motivations, it might happen that motivations change while the animal is moving at which point it reverses direction and moves back to the source of the previous bout. We assume that the ongoing performance of the animal may be interrupted whilst eating food, drinking water or moving between food and water sources. The measure that quantifies the overall performance of the animal is the expectation value of the penalty, calculated based on the geometric distribution introduced in Eq. (1). The expected penalty is given as [41]

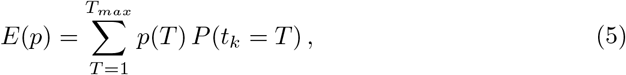

where *p*(*T*) denotes the penalty if nutritional intake stops at time *T* and *P* (*t_k_* = *T*) is the probability representing the geometric distribution as introduced in Eq (1). In the following analysis, we assume an interruption probability of λ = 0.05, yielding a maximum bout time of *T_max_* = 91, where we have added one integer time step to account for possible numerical inaccuracies so that at least 99% of the geometric distribution is considered.

In our study, we focused on the dependency of the expected penalty in Eq (5) on model parameters of the neural circuit. In particular, we numerically simulated the ongoing decision-making task by varying the cross-inhibition strength *β* and the excitation-over-inhibition ratio (E/I-ratio) defined as *r* = *α/β*. We then calculated the expected penalty to find parameter values that yield the best performance (lowest value of the expected penalty). Additionally, we determined the effect of varying the switching cost *τ* on the feeding and drinking behaviour and compared results for different initial nutritional deficits, *d*_1_ and *d*_2_, and decay parameter *γ* representing the speed by which deficits are reduced during feeding or drinking bouts. Our results are explained using bifurcation analysis. Throughout the main paper and in S1 Text we make use of the numerical continuation tool *MatCont* [70, 71] to obtain the bifurcation points when the system undergoes transitions between different dynamic regimes.

## Acknowledgments

The authors thank Philip Holmes, Jonathan Cohen, Naomi Leonard and Sebastian Musslick (all at Princeton University, NJ, US), and Benoît Girard (CNRS, Sorbonne Université, Paris, France) for fruitful discussions of the initial results of this study.

## Supporting information

**S1 Text. Supporting Information of ‘Inhibition and excitation shape activity selection: effect of oscillations in a decision-making circuit’** This document provides mathematical details and model analysis that further support our findings presented and discussed in the main paper.

## Author Contributions

T.B. and J.A.R.M. conceived of the study. T.B. designed the study, generated the data, performed the bifurcation analysis, plotted the results, and drafted the manuscript. All authors contributed to the interpretation of the results, edited the manuscript, and gave final approval for submission.

