## Supplementary material for "Inhibition and excitation shape activity selection: effect of oscillations in a decision-making circuit": S1 Text

### S1 Text: Supporting Information of 'Inhibition and excitation shape activity selection: effect of oscillations in a decision-making circuit'

Thomas Bose[1*], Andreagiovanni Reina[1], and James A. R. Marshall[1]

[1]Department of Computer Science, University of Sheffield, UK
*

July 13, 2018

#### 1 Magnitude-sensitivity

Recently, it has been observed that participants, humans in perceptual decision-making [1, 2] and monkeys in value-based decision-making [2], respond faster with increasing input magnitudes or reward values, respectively. These observations remind of Piérons law – a decay in processing time with increasing stimulus intensity – which has previously been shown to hold for binary choice experiments [3]. As discussed in [4, 5], this feature is also present in a value-sensitive model of the nest-site selection of honeybees [6, 7], which includes stop-signalling in the form of cross-inhibition [8] and, moreover, also shows other parallels with human perceptual decision-making [9].

In what follows we demonstrate that the interneuronal inhibition model employed in the present study (see Eq (4) in the main paper) is sensitive to momentary input magnitudes. Without loss of generality, we consider a single decision that resembles a two-alternative forced choice task: the animal has to decide to feed or to drink (without performing the action chosen). We numerically integrate Eq (4) presented in the main paper until either $x_1$ (motivation to feed) or $x_2$ (motivation to drink) crosses a predefined threshold $z$ for the first time. At this point the response is recorded. The plot in Fig S1 clearly shows magnitude-sensitivity, which has been proposed as a feature of effective ecological decision-making [4, 6]. Each data point represents a different condition and is based on $10^4$ simulations of the system in Eq (4), which was defined in the main paper. The graphic shows the mean decision time $\langle DT \rangle$ depending on the mean deficit $d_m = (d_1 + d_2)/2$, where $d_1$ and $d_2$ represent food and water deficits, respectively (for more details see *Methods* section in the main paper). We consider three different cases; $\Delta d = 0$, $\Delta d = 2$ and $\rho_d = 2$, where we have defined the deficit difference $\Delta d = d_1 - d_2$ and the deficit ratio $\rho_d = d_1/d_2$. As illustrated in Fig S1, we find in all of the three different cases that mean decision time decreases with increasing deficits, i.e. higher deficits trigger faster decisions. This may be understood as follows. For equal deficits the animal should decide quickly if deficits are large. The rationale behind this result is that high deficits need to be balanced as fast as possible and if deficits as well as the nutritional requirements regarding both nutrients are equal then choosing either of the available alternatives will lead to equivalent physiological states. If, on the other hand, deficits are equal and low, then allowing more time to make a decision makes it more likely that higher quality nutritional items may be detected, or that another alternative may be discovered, which possibly offers both required nutrients in one and the same food item.

The two other curves labelled $\Delta d = 2$ and $\rho_d = 2$ describe asymmetric decision problems, where food deficit $d_1$ is larger than water deficit $d_2$. In that case, a reasoning similar to the equal

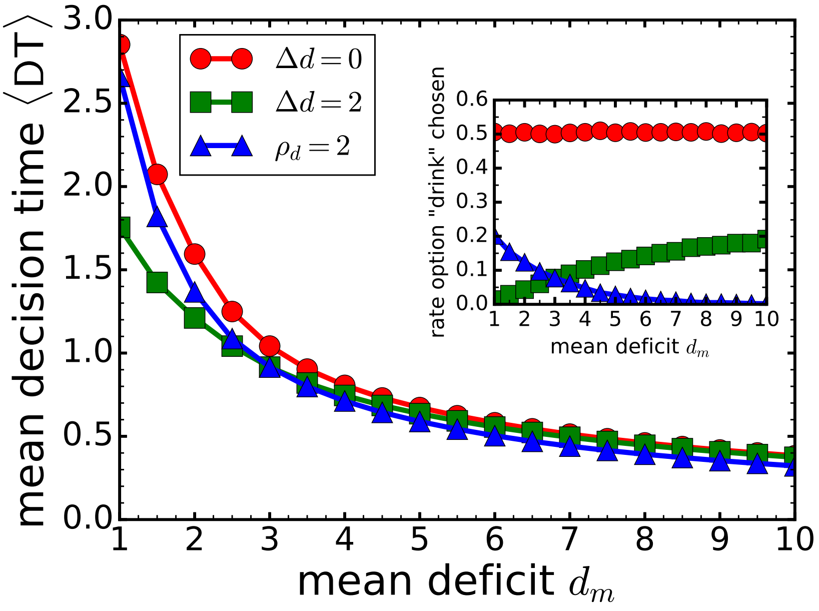


Figure S1: Plot of the mean decision time versus mean deficit. Eq (4) from the main paper is numerically integrated until threshold $z$ is crossed, i.e. either $x_1 \geq z$ or $x_2 \geq z$, whichever reaches the threshold first. Each data point is obtained from averaging over $10^4$ simulations. The inset shows the relative frequency of choosing 'to drink' – the lower-deficit option if $\Delta d > 2$ or $\rho_d = 2$. In case $\Delta d = 0$ both deficits are equal. Parameters: $z = 0.5$, $\sigma = 0.1$, $q = 0.1$, $\beta = 2$, $r = 2$, $k = 0.8$, $k_{inh} = 0.8$, $w = 3$, $g_e = 10 = g_i$, $b_e = 0.5 = b_i$.

deficit case may be applied. For larger deficits faster decisions are beneficial as the animal wants to start reducing the deficit as soon as possible. In contrast, lower deficits allow the animal to integrate more evidence before deciding in favour of one of the alternatives.

In addition, the inset in Fig S1 shows that with growing mean deficit $d_m$ but constant $\Delta d > 0$ the animal chooses drinking (i.e. the lower-deficit option) more often. This is in accordance with Weber's law of just-noticeable differences in perceptual decision-making, which describes the relation between the necessary difference in a sensory stimulus compared with a background signal, such that the stimulus can be differentiated from the background. This pattern has also been observed in mathematical analyses of honeybee nest-site selection [6, 7, 9]. In our terms this means, that the larger the mean deficit $d_m$ the harder it is for the animal to discriminate between $d_1$ and $d_2$ when the difference $d_1 - d_2$ is maintained. Regarding the relative frequency of choosing the drinking option for the $\rho_d = 2$ case, the inverse is true – choosing to drink becomes more frequent with growing $d_m$. This is also not surprising because the difference between $d_1$ and $d_2$ becomes larger for increasing $d_m$ when fixing the ratio $\rho_d$. However, at $d_m = 3$ the results for $\Delta d = 2$ and $\rho_d = 2$ coincide, as in both cases $d_1 = 4$ and $d_2 = 2$. Fig S1 was obtained for a specific set of model parameters and, although we observe the same qualitative behaviour discussed above for a wide range of parameter configurations, we note that there exist parameter regimes which show qualitatively different decision states. A detailed analysis of these states is presented in Section 2 below.

We have shown in Fig S1 that the interneuronal inhibition model studied here shows a decrease in decision time with increasing deficit-magnitudes, which are the inputs to the decision-making circuit. As our observations regarding the sensitivity to input signals are in line with previous studies in a variety of decision-making systems and contexts [1–7, 10], we believe that the applicability of the model employed in the present study bears the potential to go beyond the analysis of foraging decisions.

#### 2 Characterisation of the dynamics of the interneuronal inhibition model

Here, we present a thorough bifurcation analysis, which explains the occurrence of different dynamic regimes offered by the interneuronal inhibition model in Eq (4) in the main paper. We can quantify the distinct regimes by inspecting the bifurcation diagram in Fig S2. In this diagram we

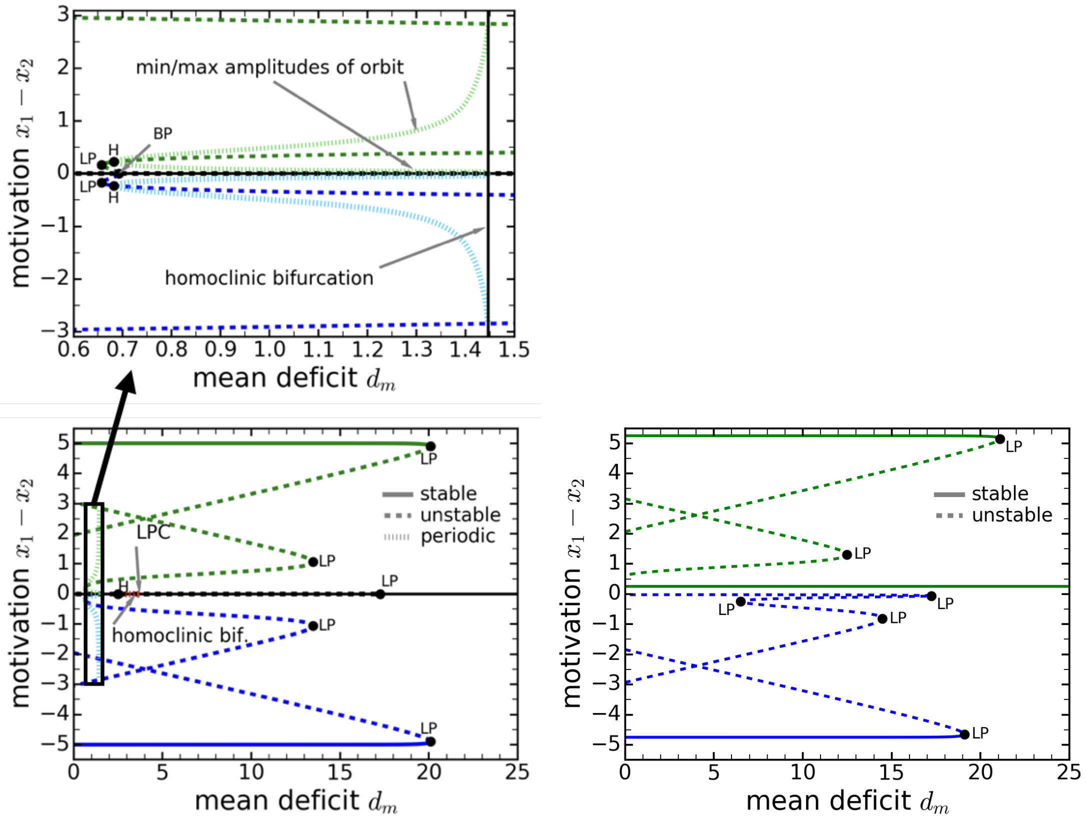


(a) equal deficits $\Delta d = 0$

(b) unequal deficits $\Delta d = 2$

Figure S2: Bifurcation plots corresponding to Fig S1, obtained with the deterministic equations in Eq (4) in the main paper ($\sigma = 0$). The values of the other parameters are: $q = 0.1$, $\beta = 2$, $r = 2$, $k = 0.8$, $k_{inh} = 0.8$, $w = 3$, $g_e = 10 = g_i$, $b_e = 0.5 = b_i$. Maximum and minimum amplitudes are plotted for the periodic solutions. The upper panel in (a) shows the enlarged area indicated in the lower panel. Abbreviations: LP: limit point, BP: branch point, H: Hopf bifurcation point, LPC: limit point of cycles.

chose the mean deficit $d_m$ as bifurcation parameter and plot the motivation difference $\Delta x = x_1 - x_2$ depending on varying $d_m$, which correspond to the $\Delta d = 0$ and $\Delta d = 2$ curves in Fig S1. The equal alternatives case depicted in Fig S2a shows stable fixed points over a wide range of deficit values. We observe that for $d_m \in [0, 20.1]$ high-difference solutions and $\Delta x = 0$ solutions coexist. For $d_m > 20.1$ the stable attractor persists, which describes a decision-deadlock state ($\Delta x = 0$). The other two vanish via saddle-node bifurcations (rightmost limit points). In addition, for $d_m < 20.1$ we find three more limit points as indicated in Fig S2a. We also observe three periodic solutions that emerge via Hopf bifurcations. The Hopf bifurcation point at $d_m \approx 2.5$ is characterised by $\Delta x = 0$, i.e. maximum and minimum amplitudes of that periodic cycle are equal and zero. However, the remaining two Hopf bifurcations occurring at $d_m \approx 0.7$, show nonzero amplitudes when subtracting $x_2$ from $x_1$. This is emphasized in the upper panel in Fig S2a which shows an enlarged view of the periodic oscillations. Although only the periodic orbit with $\Delta x = 0$ exhibits a limit point of cycles, where stable (smaller amplitudes) and unstable (larger amplitude) oscillations meet and annihilate each other in a fold bifurcation of periodic orbits, all limit cycles eventually vanish via a homoclinic bifurcation. This is where a periodic orbit and a saddle point meet and an unstable manifold of the saddle blows up the amplitude of the limit cycle, leading to a strong increases of the periods of the orbits.

In contrast, in Fig S2b periodic solutions in the range of biologically accessible positive deficits $d_m$ are lost when introducing the deficit difference $\Delta d = 2$. In addition to several branches that are unstable (here we use the term unstable if at least one of the three eigenvalues of the fixed point has a positive real part), we observe three stable equilibria: one equilibrium with $\Delta x < 0$ and two equilibria with $\Delta x > 0$. Comparing this with Fig S2a we see that those curves are now upshifted because we have assumed a nonzero deficit difference. However, we still observe that for large values of $d_m$ there is only one stable attractor, where $\Delta x > 0$ but small. Just as in the symmetric case ($\Delta d = 0$) the two rightmost limit points occurring at $d_m = 19.1$ and $d_m = 17.2$ determine the transition into a decision deadlock state. Due to the asymmetry of the decision problem ($\Delta d > 0$) this transition takes place in two steps: the tristable regime becomes bistable when increasing $d_m$ above 17.2 and bistability is lost for $d_m > 19.1$, where only one stable solution remains. Hence, considering the dependence of the decision outcome on nutritional deficits and deficit differences as above clearly shows that the physiological state of the animal may strongly affect the final decision state of the animal. However, to account for varying deficits the neural circuit underlying the decision-making process should be adaptive. In what follows, adaptivity is studied by varying the inhibition strength $\beta$, which we use as the critical parameter in the bifurcation diagram in Fig S2. The importance of inhibitory mechanisms in the brain is well known (e.g. see [11]), and has been emphasized in perceptual decisions in human and primate decision makers (e.g. see [1, 12–15]), and in value based decisions in house hunting honeybees [6–8]. Thus, we expect the inhibition parameter $\beta$ to play a key role in our model, too.

The equal deficits case ($\Delta d = 0$) is illustrated in Fig S3a. When varying the values of the bifurcation parameter $\beta$, we see a transition from a monostable to a tristable regime, which are separated by two limit points at $\beta \approx 0.34$. This means that we observe a decision-deadlock state ($\Delta x = x_1 - x_2 = 0$) for inhibition strengths $\beta < 0.34$ and possible deadlock-breaking for $\beta > 0.34$, as there are now two additional stable fixed points. Regarding the latter, we find additional bifurcations that further divide the bifurcation diagram. We also observe two branch points – one occurring at $\beta \approx 0.34$, i.e. at approximately the same inhibition strength where the limit points are observed, and another one at $\beta \approx 0.79$. From the latter we see two stable branches emerging that lose stability at Hopf bifurcation points at $\beta \approx 1.05$, where two stable limit cycles are born. When increasing $\beta$ further we detect a limit point of cyclces as indicated in Fig S3a describing fold bifurcations of the two periodic orbits. The periodic solutions are lost via homoclinic bifurcations. The upper panel in Fig S3a shows a magnified view on the limit cycles. Furthermore, there is another Hopf bifurcation found at $\beta \approx 1.85$. Also this family of periodic solutions undergoes a limit point bifurcation of cycles and a homoclinic bifurcation at which point the periodic solutions vanish.

Making the comparison between equal deficits ($\Delta d = 0$) and unequal deficits ($\Delta d = 2$), several changes can be recognised in Fig S3b. Introducing a nonzero $\Delta d$ lifts the degeneracy which was observed in Fig S3a. This means that curves, which overlay in the $\Delta d = 0$ case along the $\Delta x = 0$ line, are separated for $\Delta d = 2$. We also see that one limit point and one branch point have moved together, and the $\beta$-values are numerically indistinguishable. The other branch point observed at $\beta \approx 0.79$ in Fig S3a has become disconnected and is now a limit point. If $\Delta_d = 2$ we observe monostabilty for $0 \leq \beta \leq 0.16$, bistability for $0.16 < \beta \leq 0.47$, and quadristability (four stable fixed points) for $0.47 \leq \beta \leq 0.72$, and tristability for $\beta > 0.72$. In addition, in the small range $0.72 \leq \beta \leq 0.76$, we also detect a limit cycle, which however does not exist outside this interval. Corresponding limit cycle solutions are shown in an enlarged view in the upper panel of Fig S3b. To better resolve all bifurcation events we confined the area depicted in Fig S3 to $|\Delta x| \leq 5$ and $0 \leq \beta \leq 2.5$. We also increased the inhibition strength to $\beta$=20 and did not observe further qualitative changes for $\beta > 0.76$. This means the area of tristability persists, where one stable

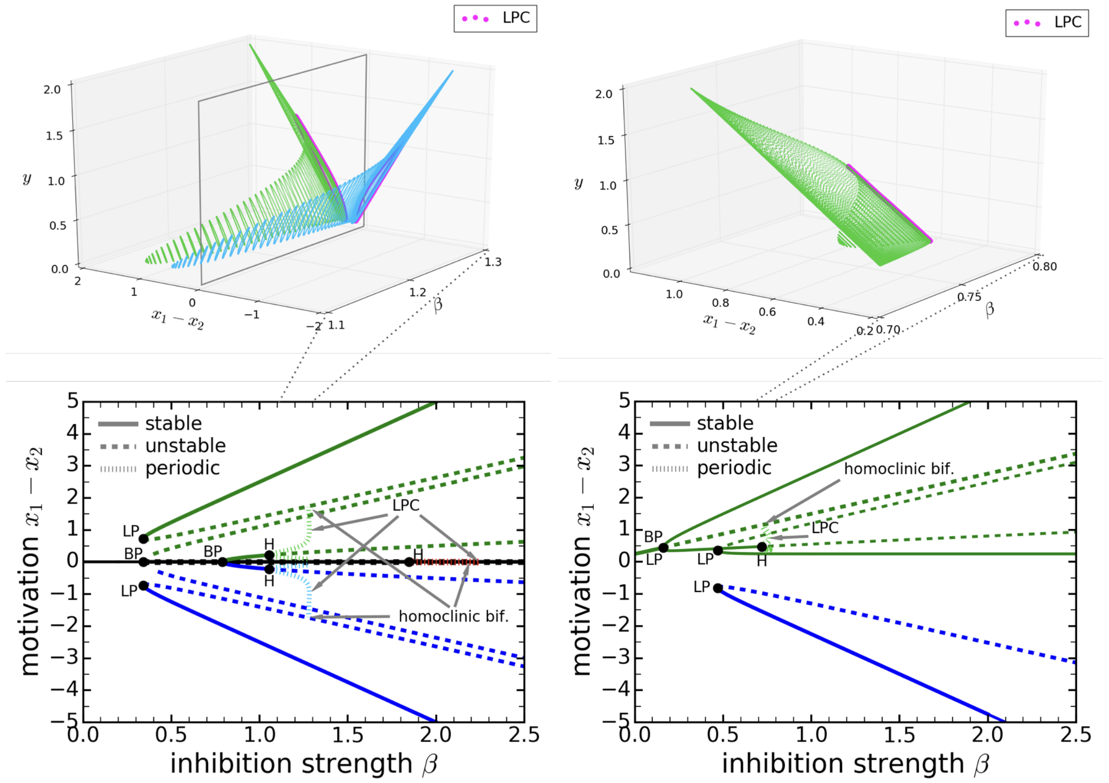


(a) equal deficits $\Delta d = 0$

(b) unequal deficits $\Delta d = 2$

Figure S3: Bifurcation plots for equal (a) and unequal (b) deficits corresponding to $d_m = 3$ in Fig S1, obtained with the deterministic set of equations in Eq (4) in the main paper ($\sigma = 0$). The critical parameter is the inhibition strength $\beta$. Maximum and minimum amplitudes are plotted for the periodic solutions. The upper panels in (a) and (b) show the periodic solutions in an increased view. Parameters: $q = 0.1$, $r = 2$, $k = 0.8$, $k_{inh} = 0.8$, $w = 3$, $g_e = 10 = g_i$, $b_e = 0.5 = b_i$. Abbreviations: LP: limit point, BP: branch point, H: Hopf bifurcation point, LPC: limit point of cycles.

equilibrium is characterised by a low positive motivation difference and the other two by large motivation differences both positive and negative. Having negative differences as available stable solutions has the consequence that the nutritional alternative in which the animal has the lower deficit might be chosen. Compared with the higher-deficit alternative, the lower-deficit alternative will, however, be less likely to be chosen as the stable branch $\Delta x < 0$ is disconnected from the other stable solutions. Assuming that, starting from $\beta \approx 0$, the animal may increase the inhibition parameter over time, then it is more probable that the solution stays on one of the stable branches with $\Delta x > 0$. Increasing inhibition strengths over time has been suggested to be operative in the nest-site selection of honeybees [6–8], for example.

#### 3 Evidence that cross-inhibition model offers no periodic solution

Here we present evidence that closed orbits do not occur in the cross-inhibition model (equivalently termed mutual inhibition model) which we express as

$$\begin{aligned} \frac{dx_1}{dt} &= -kx_1 + \alpha f_e(x_1) - \beta f_i(x_2) + I_1(d_1)\,, \\ \frac{dx_2}{dt} &= -kx_2 + \alpha f_e(x_2) - \beta f_i(x_1) + I_2(d_2)\,, \end{aligned} \tag{S1}$$

where $f_{e,i}$ are sigmoidal functions defined in Eq (3) in the main paper (with the same shape as the sigmoidal function in Eq (S4) given below) and all the model parameters have the same meaning as before. Important differences between mutual and interneuronal inhibition models are that the latter is three-dimensional whilst the former is a two-dimensional model. Moreover, the arguments of the inhibition functions $f_i$ are now the motivations $x_1$ and $x_2$ instead of the interneuronal activity level denoted $y$ before. This implementation of the cross-inhibition motif is comparable with previous models of decision making (cf. [16–18]) and it is on the same mathematical footing compared with interneuronal inhibition motif studied in Eq (4) in the main paper.

To give evidence that the system in Eq (S1) does not offer closed periodic orbits, we make use of the Dulac-Bendixson criterion, e.g. see [19]. We consider the $C^1$ function $\phi(x_1,x_2)$ and calculate

$$L(x_1,x_2) = \frac{\partial\,[\phi(x_1,x_2)\,\psi_1(x_1,x_2)]}{\partial x_1} + \frac{\partial\,[\phi(x_1,x_2)\,\psi_2(x_1,x_2)]}{\partial x_2} \tag{S2}$$

where $\psi_1$ and $\psi_2$ represent the right-hand sides of Eq (S1), i.e. $\psi_1 = \dot{x}_1$ and $\psi_2 = \dot{x}_2$. First, we consider the case without leak, that is $k = 0$. Letting $\phi(x_1,x_2) = 2/(\alpha g_e)$ we find that the expression in Eq (S2) yields

$$L(x_1,x_2) = 2\sum_{j=1}^{2} \frac{\exp[-g_e\,(x_j - b_e)]}{(1+\exp[-g_e\,(x_j - b_e)])^2}\,, \tag{S3}$$

for the mutual inhibition model. The Dulac-Bendixson criterion states that if $L \neq 0$ does not change sign then there are no limit cycles or homoclinic connections occurring in the system. Inspecting Eq (S3) we find that $L$ has its maximum value at $x_1 = b_e = x_2$, i.e. $L_{max} = L(x_1 = b_e, x_2 = b_e) = 1$. For all other values of $x_1$ and $x_2$ we have $0 < L < 1$. In particular, letting $x_{1,2} \to \pm\infty$ then $L \to 0$ asymptotically. Hence, the mutual inhibition model satisfies the Dulac-Bendixson criterion if $k = 0$. In case $k > 0$ we did not find a simple expression for the Dulac-function $\phi(x_1,x_2)$. Therefore, we present a heuristic argument. It is well known from molecular regulatory networks that in order to observe periodic solutions we need a coupling in the system of differential equations containing both positive and negative feedback, or a third population mediating negative feedback [20]. The latter applies to the interneuronal inhibition model in Eq (4) in the main paper. As we have no cross-excitation mechanism in the mutual inhibition model there is no opposing force to the cross-inhibition. We suppose that a nonzero leak parameter $k$ does not change this situation, as it only models the loss of information over time and should therefore not enable the system to enter an oscillatory state leading to closed orbits. In addition, we also performed a numerical study of the system in Eq (S1) and did not observe closed periodic orbits. Further to this, no other work studying the cross-inhibition motif ([16–18]) reported their existence.

Taking together the mathematical proof for $k = 0$ and our heuristic argumentation for $k > 0$ we are confident that our assumption about the non-existence of periodic orbits in the mutual inhibition model is valid. Our study of the interneuronal inhibition motif is therefore well-motivated as it allows for the occurrence of periodic limit cycles as an additional feature.

### 4 Validity of approximating sigmoidal function with linear function for deficit-motivation coupling

Sigmoidal functions used in this study have the general shape given as

$$f(\xi) = \frac{1}{1+\exp[-g\,(\xi - b)]}\,, \tag{S4}$$

where $g$ and $b$ denote the gain and the inflection point of $f$. We can approximate the function $f$ around the inflection point $b$ by a Taylor expansion at $\xi = b$. Considering terms up to first order, this yields

$$f(\xi) \simeq \frac{1}{2} + \frac{g}{4}(\xi - b) =: f_{lin}(\xi)\,, \tag{S5}$$

where we have introduced the linear approximation of $f$ denoted $f_{lin}$. Under the assumption that $g = 2/b$ and introducing the deficit $d = \xi$, we may express the linear approximation of $f(\xi \to d)$ as $f_{lin}(d) = d/(2b)$. The input functions, $I_1(d_1)$ and $I_2(d_2)$, in Eq (2) in the main paper may be obtained by equating $I_j = \kappa f_{lin}(d_j)$, where $\kappa$ is a constant. Introducing further $q = \kappa/(2b)$ yields the expressions for the input functions in Eq (2). This means that the linear coupling applied in the interneuronal inhibition model in Eq (4) can be understood as an approximation of a nonlinear relationship. Although the sigmoidal dependence is just one possibility to introduce the coupling of nutritional deficits and motivations, it is a reasonable assumption. For low deficits there is only a low incentive for the animal to forage and for high deficits there is a much larger drive to feed and drink. However, as all resources inside the body are limited there should also be an upper limit on the behaviour driven by deficits. Hence, a deficit-depending function that drives foraging decisions should saturate at some deficit value, as described by the sigmoidal function in Eq (S4). Approximating the sigmoidal function by a linear one in our model should be valid for moderate deficits. This is illustrated in Fig S4. As shown in this graph, assuming deficits

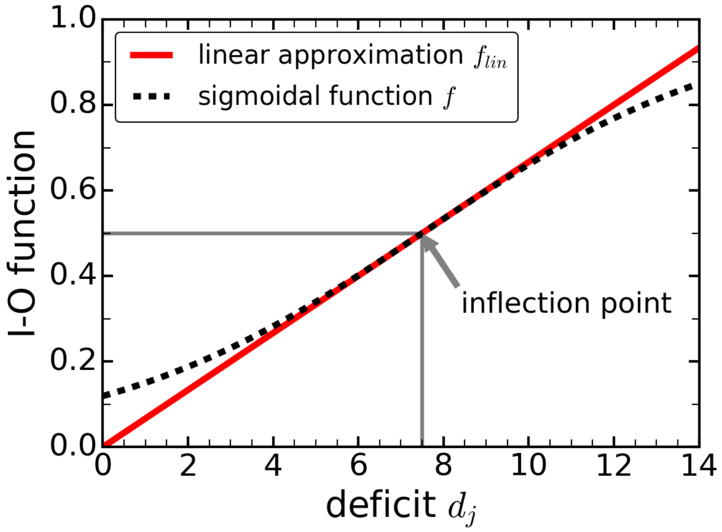


Figure S4: Linear approximation of sigmoidal function with parameters $g = 2/b$ and $b = 7.5$.

$d_j \in [2, 10]$ confirms that the linear approximation should work reasonably well. In our study, deficit values almost exclusively range in that interval, which makes the linear coupling in Eq (4) in the main paper a valid model assumption.

### 5 Linearisation of interneuronal inhibition model

In the following we derive the equations of motion which underly Figs 4D-F in the main paper and Fig S8b below. To find the corresponding system of equations we make use of the linear approximation derived in the previous section, i.e. approximating Eq (S4) by Eq (S5). Applying this expansion to the sigmoidal functions $f_e$ and $f_i$ in Eq (4) in the main paper and collecting terms of equal order, we get

$$\begin{aligned}
dx_1 &= \left[-\left(k-\frac{r\beta g_e}{4}\right)x_1 - \frac{\beta g_i}{4}y + q d_1 + \varphi\right] dt + \sigma\, dW_1\,,\\
dx_2 &= \left[-\left(k-\frac{r\beta g_e}{4}\right)x_2 - \frac{\beta g_i}{4}y + q d_2 + \varphi\right] dt + \sigma\, dW_2\,,\\
dy &= \left[-k_{inh}\, y + w\left(1-\frac{g_e b_e}{2} + \frac{g_e}{4}(x_1+x_2)\right)\right] dt\,,\\
\varphi &= \frac{1}{2}\beta\left[r\left(1-\frac{g_e b_e}{2}\right) - \left(1-\frac{g_i b_i}{2}\right)\right],
\end{aligned} \tag{S6}$$

as the linearised version of the interneuronal inhibition model, which has been used to obtain the results shown in Figs 4D-F in the main paper and Fig S8b below. Here, like in the main paper, we have introduced the E/I-ratio $r = \alpha/\beta$. All model parameters have the same meaning as in Eq (4) in the main paper.

### 6 Equivalence with previous model of perceptual and nutritional decision making

In this section we relate the approximated linear interneuronal inhibition motif in Eq (S6) to previous models. This can be achieved under the assumption that the decay of inhibitory activity is fast compared with the decay of excitatory activity [13, 14], where excitatory activity is given by the motivational state in our model. Mathematically this means in our terms that $k_{inh} \gg (k - r\beta g_e/4)$. Then, considering the motivational time scale, the activity of the interneuronal unit operates at its steady state ($dy/dt = 0$) and may be expressed as

$$y = \frac{w}{k_{inh}}\left(1-\frac{g_e b_e}{2} + \frac{g_e}{4}(x_1+x_2)\right). \tag{S7}$$

Inserting Eq (S7) in Eq (S6), and setting $\sigma = 0$ (i.e. assuming a deterministic system) we obtain

$$\begin{aligned}
\frac{dx_1}{dt} &= \tilde{c}_1 + c_2 d_1 + c_3 x_1 + c_4 x_2\,,\\
\frac{dx_2}{dt} &= \tilde{c}_1 + c_2 d_2 + c_3 x_2 + c_4 x_1\,,
\end{aligned} \tag{S8}$$

where the constants $\tilde{c}_1$, $c_2$, $c_3$ and $c_4$ are expressed as

$$\begin{aligned}
\tilde{c}_1 &= \varphi + \frac{w}{k_{inh}}\left(1-\frac{g_e b_e}{2}\right), & c_2 &= q\,,\\
c_3 &= -\left(k + \frac{\beta g_e}{4}\left[\frac{w g_i}{4k_{inh}} - r\right]\right), & c_4 &= -\frac{w\beta g_e g_i}{16 k_{inh}}\,,
\end{aligned} \tag{S9}$$

and the expression for $\varphi$ is given in Eq (S6).

The model in Eq (S8) closely resembles the one studied in [21]. The only difference is the form of the $\tilde{c}_1$ term. Here, $\tilde{c}_1$ is a constant given by the model parameters of the neural circuit. In [21] this is replaced by $\tilde{c}_1 \to c_1 \dot{d}_j$ $(j = 1,2)$, where $c_1$ is a constant and the $\dot{d}_j$ represent the time-derivative of the deficits. Although $\dot{d}_j$ is given by a constant in [21] as well as in the present paper (deficit decay parameter $\gamma$), deficits are only reduced whilst the animal is consuming nutrients. This means $\dot{d}_j = 0$ $(j = 1,2)$ when the animal is located somewhere between water and food sources and $\dot{d}_j = -\gamma$ $(j = 1,2)$ if the model animal feeds at the food source or drinks at the water source, respectively. Therefore, exact equivalence is achieved when $c_1 = 0$ in [21] and $\tilde{c}_1 = 0$ in the present paper. The condition $\tilde{c}_1 = 0$ is obtained for the special case when $g_e b_e = 2 = g_i b_i$.

However, starting from the nonlinear interneuronal inhibition model in Eq (4) in the main paper and arriving at the linear two-dimensional system in Eq (S8) (via the linearised interneuronal inhibition model in Eq (S6)) makes it possible to express the model constants in the approximated cross-inhibition model in (S8) by neurobiological parameters, as demonstrated in Eq (S9).

#### 7 Comparison of deficit reduction for different switching costs

In Fig S5 we show a comparison of the deficit reduction in the ongoing decision making task of the model animal in dependence on the physical distance between food and water source given as switching cost $\tau$. In agreement with other work [21, 22] the animal performs better when the

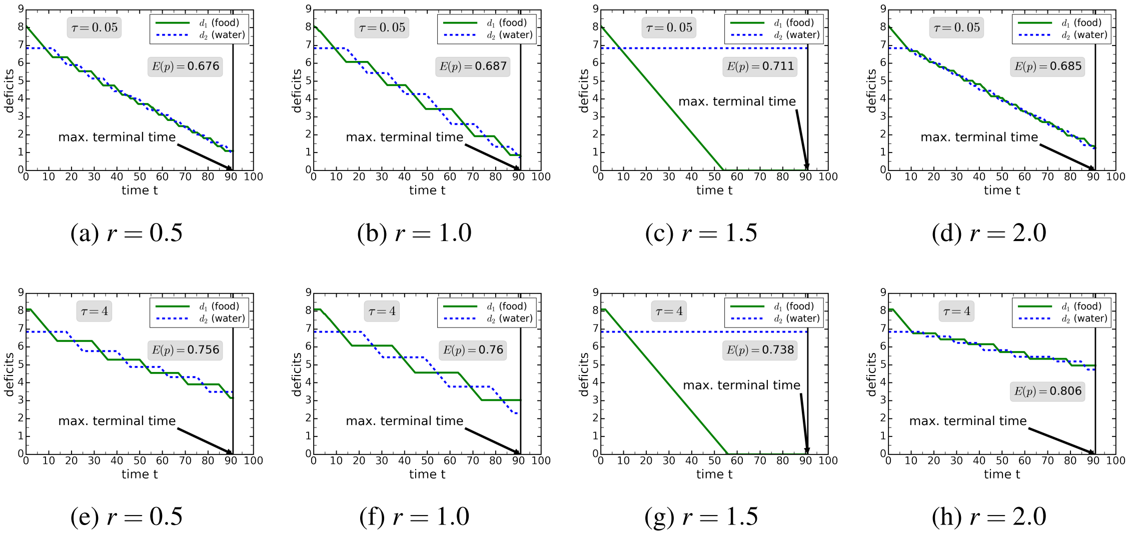


(a) $r = 0.5$ (b) $r = 1.0$ (c) $r = 1.5$ (d) $r = 2.0$

(e) $r = 0.5$ (f) $r = 1.0$ (g) $r = 1.5$ (h) $r = 2.0$

Figure S5: Effect of switching cost $\tau$ and E/I-ration $r$ on deficit reduction. We chose a symmetric starting point at $\tau/2$. The expected penalties, $E(p)$, and switching costs (travel time between food and water source), $\tau$, are given in each plot. Lower penalty values mean better performance of the model animal. Parameters: $d_m(t = 0) = 7.47$, $\Delta d(t = 0) = 1.25$, $\tau = 4$, $\beta = 3$, $\gamma = 0.15$, $q = 0.1$, $k = 0.8$, $k_{inh} = 0.8$, $w = 3$, $g_e = 10 = g_i$, $b_e = 0.5 = b_i$, and $\sigma = 0.01$.

switching cost between feeding and drinking is lower, compare Figs S5a-S5d with their counterparts in Figs S5e-S5h. The animal also performs better when it alternates between feeding and drinking, if $\tau$ is sufficiently low. However, if the opposite applies and the switching cost is significantly higher, then animals performing exclusively one activity could improve there performance at the end of the ongoing foraging task. This behaviour can be achieved by modulating the E/I-ratio accordingly. Figures S5e-S5h illustrate this result. The animal performing only one activity achieves the best performance for $\tau = 4$ (see Fig S5g). As discussed in the main paper, this observation is a direct consequence of the nonlinearity of the underlying Eq (4), and is of course only reasonable in the short-term, to which our study refers. In contrast, over longer periods of time the animal needs to perform both activities to survive.

#### 8 Dependence of expected penalty on switching cost $\tau$

The switching cost denoted $\tau$ in our study measures the time the model animal needs to overcome the distance between food and water source. During that time the animal cannot reduce any of the deficits, therefore switching and travelling is costly. How this cost influences the performance of the animal is shown in the following. In particular, we have assumed initial deficits $d_1(t = 0) = 7.5 = d_2(t = 0)$ and compared the expected penalties for five different values of $\tau$, i.e. $\tau = 2, 4, 8, 16$ and $32$. The corresponding results are depicted in Fig S6. We can recognise that,

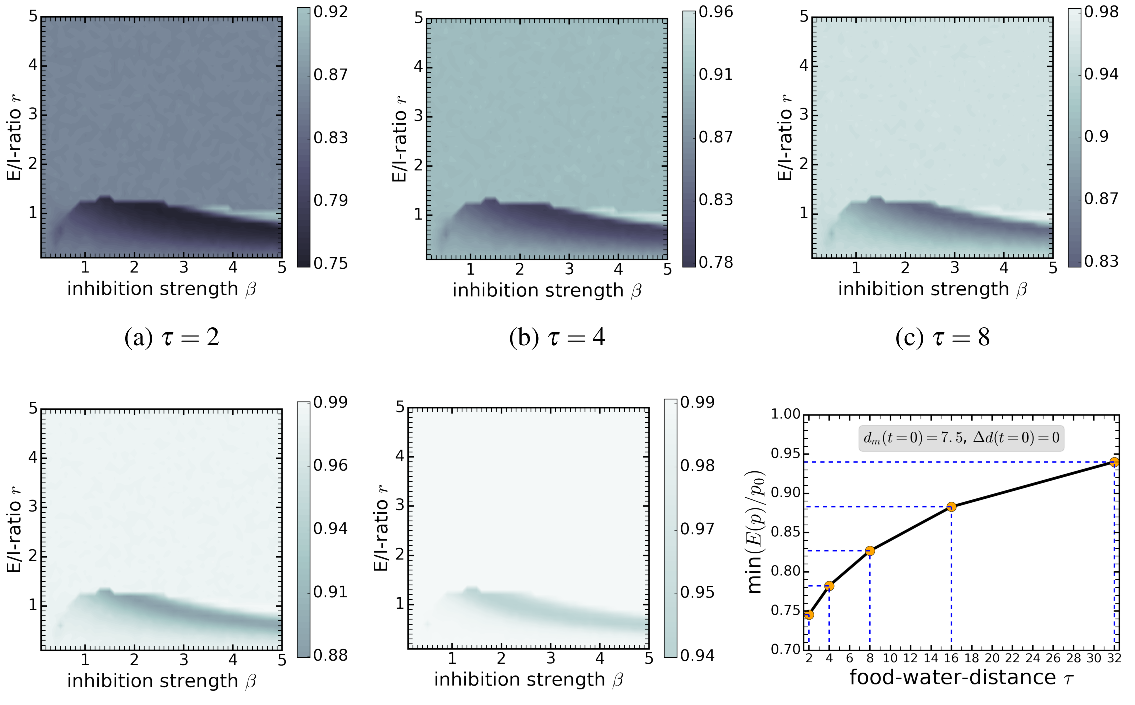


(a) $\tau = 2$ (b) $\tau = 4$ (c) $\tau = 8$

(d) $\tau = 16$ (e) $\tau = 32$ (f) minimum penalty values

Figure S6: Dependence of expected penalty on distance between food and water sources ($\tau$). We chose a symmetric starting point at $\tau/2$ in all plots. Areas characterised by the lowest penalty values mirror the best performance of the model animal. Parameters: $d_m(t=0) = 7.5$, $\Delta d(t = 0) = 0$, $\gamma = 0.15$, $q = 0.1$, $k = 0.8$, $k_{inh} = 0.8$, $w = 3$, $g_e = 10 = g_i$, $b_e = 0.5 = b_i$, and $\sigma = 0.01$.

although the shape of the penalty landscape in Figs S6a-S6e remains very similar under variation of $\tau$, the whole process becomes less effective. A numerical comparison of the minimum values of the normalised expected penalties $\min(E(p)/p_0)$ after terminal time $T_{max}$, $p_0 = d_1^2(t=0) + d_2^2(t=0)$ being the initial penalty value, is shown in Fig S6f. The shape of the curve in this diagram confirms that success decreases with increasing switching cost. On the one hand this was to be expected, but on the other hand observing this behaviour provides evidence for the correctness the computational implementation of our model, and it also demonstrates that an increase in $\tau$ leads a nonlinear relationship between expected penalty and switching cost.

### 9 Further details on the dependence of expected penalty on initial deficits

In order to understand how different initial deficits affect the behaviour of the model animal, we analysed performance plots, i.e. expected penalty depending on $r$ and $\beta$, for different initial deficits $d_j(t=0) = 5, 7.5$ and $10$, $j = 1, 2$. This is illustrated in Fig S7. Inspecting the diagrams in Figs S7a-S7c, it becomes obvious that the shape of the performance plot changes when altering the initial deficit $d_m(t=0)$. Increasing the initial deficit is accompanied by two effects. First, the area of improved performance becomes smaller, i.e. for higher initial deficits (Fig S7c) improved performance is only observed for higher inhibition strengths $\beta$ and lower E/I-ratios $r$, compared with lower initial deficits (Fig S7a). Second, the normalised penalty is increased when going to higher initial deficits. Here, different values of $p_0$ where used in Figs S7a-S7c, corresponding to the different initial deficits. This effect is emphasised again in Fig S7d, where we plotted the minimum value of the normalised penalties Figs S7a-S7c. However, looking at the inset in Fig S7d reveals that the absolute difference of penalty values $\Delta E(p) = p_0 - \min(E(p))$ increases

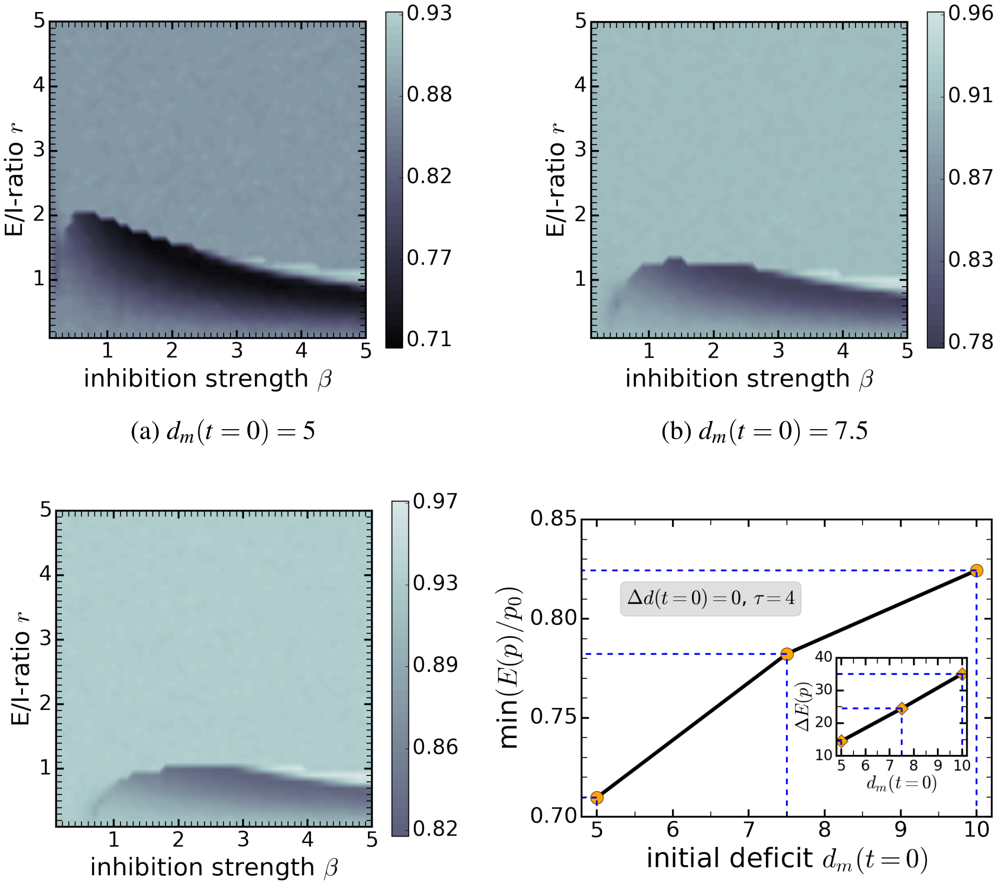


(a) $d_m(t=0)=5$

(b) $d_m(t=0)=7.5$

(c) $d_m(t=0)=10$

(d) minimum penalty values

Figure S7: Dependence of expected penalty on initial deficits $d_m(t=0)$. We chose a symmetric starting point at $\tau/2$. Areas characterised by the lowest penalty values mirror the best performance of the model animal. Parameters: $\Delta d(t=0)=0$, $\tau=4$, $\gamma=0.15$, $q=0.1$, $k=0.8$, $k_{inh}=0.8$, $w=3$, $g_e=10=g_i$, $b_e=0.5=b_i$, and $\sigma=0.01$.

with increasing initial deficits. This means that in absolute terms the expected penalty is more reduced for higher initial deficits than it is for lower initial deficits, given the same duration of the decision-making task. This reflects the feature of the interneuronal inhibition model discussed in Section 1: higher magnitudes (deficits) evoke faster decisions. As the deficit decay constant $\gamma$ for food and water intake is identical for the different cases in Fig S7, we conjecture that it is the magnitude-sensitivity of the interneuronal inhibition model which is responsible for this effect.

#### 10 Dependence of expected penalty on deficit decay parameter $\gamma$

Additionally, we depict the dependencies of the expected penalty on the deficit decay constant $\gamma$ in Fig S8. We compare results obtained for the nonlinear model (see Eq (4) in the main paper) in Fig S8a with results for a linearised model in Fig S8b (see Eq (S6)). This graphic highlights once more that different $r$ values also cause differences in the animal's performance. In the diagram in Fig S8a, the animal achieves the best performance for an E/I-ratio $r=1$, and performs worse for $r=0.5$, $r=1.5$ and $r=2$, i.e. for both smaller and larger values of the E/I-ratio. Although expected penalty values decrease with increasing deficit decay rate $\gamma$, the different curves do not cross each other in the range of $\gamma$-values studied.

Differences in the performance when modulating the E/I-ratio in response to different nutrient decay rates are also observable using the linear version of the interneuronal inhibition model (see

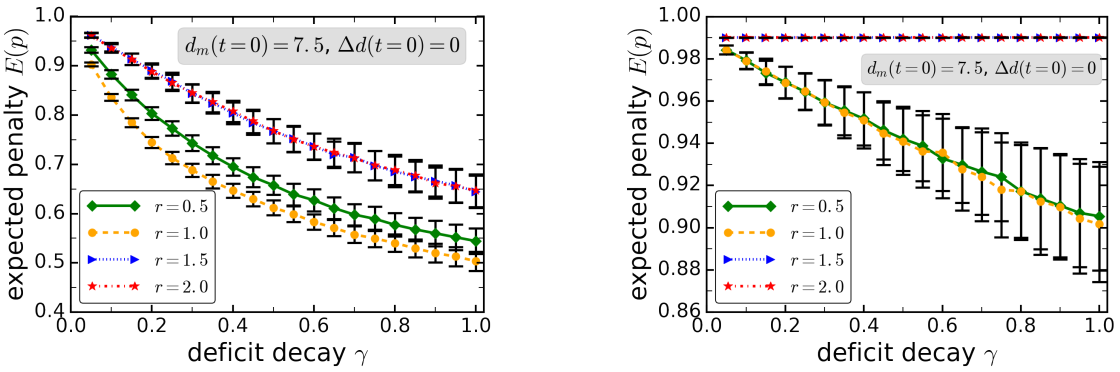


(a) nonlinear model (b) linearised model

Figure S8: Plot of expected penalties depending on the constant decay parameter $\gamma$, describing the reduction of nutritional deficits whilst the animal feeds or drinks. Curves are shown for different E/I-ratios. Error bars represent standard deviations obtained from 1000 simulations. (a) nonlinear model in Eq (4) in the main paper; (b) linearised model presented in Eq (S6). Parameters: $d_m(t=0)=7.5$, $\Delta d(t=0)=0$, $\tau=4$, $\beta=3$, $q=0.1$, $k=0.8$, $k_{inh}=0.8$, $w=3$, $g_e=10=g_i$, $b_e=0.5=b_i$, and $\sigma=0.01$.

Fig S8b), although the performance differences occur on a smaller scale (compare the numbers on the vertical axes of Figs S8a and S8b). However, it becomes obvious that using the linear model the animal performs poorly for $r=1.5$ and $r=2$. For those two values of the E/I-ratio, the animal is subject to frequent motivational switches and can therefore not perform activities chosen for a reasonable amount of time. Moreover, it may not even reach either food or water source but moves back and forth in between the two sources – a behaviour known as dithering, which is clearly disadvantageous for the animal. Although dithering may also occur in the nonlinear model, we have not observed it on this extreme scale in our study, compared with the linearised model. Hence, nonlinear models may also be a way to avoid too frequent switches when analysing animal foraging behaviour.
